# A comprehensive evolutionary scenario for the origin and neofunctionalization of the *Drosophila* speciation gene *Odysseus* (*OdsH*)

**DOI:** 10.1101/2023.05.31.542109

**Authors:** William Vilas Boas Nunes, Daniel Siqueira de Oliveira, Guilherme de Rezende Dias, Antonio Bernardo Carvalho, Ícaro Caruso Putinhon, Joice Matos Biselli, Nathalie Guegen, Abdou Akkouche, Nelly Burlet, Cristina Vieira, Claudia M. A. Carareto

## Abstract

*Odysseus* (*OdsH*) was the first gene described in *Drosophila* related to speciation and hybrid sterility. This gene was first described in the *melanogaster* subgroup and more specifically in the sterile hybrids from crosses between *D. mauritiana* and *D. simulans*. Its origin is attributed to the duplication of the gene *unc-4*, which would have occurred in the ancestor of the subgenus *Sophophora*. By using a much larger sample of *Drosophila* species, we showed that contrary to what has been previously proposed, *OdsH* origin occurred approximately 62 million years ago (Mya). Then, *OdsH* have experienced rapid neofunctionalization in male reproductive tracts, evidenced by its evolutionary rates, expression and transcription factor binding sites. Furthermore, the analysis of the OdsH peptide sequence allowed the identification of mutations in the DNA- and protein-binding domains of *D. mauritiana* that could result in incompatibility with genomes from other species. We then explored the expression of *OdsH* in the spermatocytes of *D. arizonae* and *D. mojavensis*, a pair of recently diverged sister species with incomplete reproductive isolation and expected to find the involvement of *OdsH* in hybrid sterility. Our data indicated that *OdsH* expression is not atypical in male-sterile hybrids from these species. In conclusion, we have demonstrated that the origin of *OdsH* occurred earlier than previously proposed and that its neofunctionalization in male sexual functions occurred rapidly after its origin. Our results also suggested that its role as a speciation gene, as in the *melanogaster* subgroup of species, may be restricted to this specific taxon.

## Introduction

*Odysseus* (*OdsH)* was the first speciation gene characterized in *Drosophila*, specifically between *D. mauritiana* and *D. simulans* (Ting et al. 1998). The sterility of male hybrid offspring from this cross was due to the introgression of a sequence from *D. mauritiana* encompassing *OdsH* into the *D. simulans* genome (Perez et al. 1993; Perez and Wu 1995; Ting et al. 1998). The atypical expression of *OdsH* at the apical testis region was observed in these hybrids (Sun et al. 2004), and its translated protein was described as a heterochromatin-binding transcription factor (Bayes and Malik, 2009). The origin of the *OdsH* gene is proposed to have arisen by duplication of the *unc-4* gene, a conserved gene in Metazoa located *in tandem* with *OdsH* (Ting et al. 2004).

Duplicated genes can have different outcomes, other than being lost or pseudogenized. They can be maintained with the same function as the parental genes, increasing gene expression, and are expected to be under purifying selection (Ohno 1970). In other cases, the ancestral function is divided in complementary subfunctions in the two gene copies, which is called subfunctionalization (Hughes 1994; Force et al. 1999; Soltzfus 1999). Additionally, duplicated genes can become specialized for innovative functions, leading to the evolution of new traits or characteristics. This process is called neofunctionalization (Ohno 1970).

*unc-4*, the parental gene, is associated with motor neuron and proprioceptor developmental pathways in *D. melanogaster* (Tabuchi et al. 1998; Lacin and Truman, 2016, 2019, 2020), similar to its conserved single copy orthologue, which acts on motor neuron and optical sensorial cell development in *Caenorhabditis elegans* (Miller et al. 1992; Fox et al. 2005; Marques et al. 2019). *OdsH*, the duplicated copy, is expressed in spermatocytes in species of the *melanogaster* subgroup (Ting et al. 2004; Bayes and Malik, 2009), in agreement with the out-of-testes hypothesis, which proposes that new genes tend to be expressed first in testis and then might evolve functions in other tissues (Kaessman 2010). This has been observed in the majority of *Drosophila* newly duplicated genes that evolve new functions in the testis (Assis and Bachtrog 2013; Jiang and Assis 2017; Raices et al. 2019; Su et al. 2021).

Both genes, *unc-4* and *OdsH*, encode transcription factors with homologous DNA-binding homeodomains, phylogenetically classified in the Paired-like class (Winnier et al. 1999). Proteins carrying such motifs generally contain a C-terminal octapeptide motif that was described as interacting with the Unc-37 protein in *C. elegans* (orthologous to Groucho in *Drosophila*), acting as an expression repressor of their DNA targets (Winnier et al. 1999). The octapeptide was also identified at the C-terminal ends of both Unc-4 and OdsH in *D. melanogaster* (Copley 2005). The *OdsH* homeodomain has a high amino acid substitution rate in species from the *melanogaster* subgroup, corresponding to a higher divergence between the domains from *unc-4* between *Drosophila* and evolutionarily distant species, such as *C. elegans* (Ting et al. 2004). As expected for duplicated genes, the faster evolution of the *unc-4* paralogue is associated with the acquisition of novel functions in the testis and with the speciation process (Ting et al. 1998, 2004). Several studies on ancient duplicates propose that neofunctionalization is driven by positive selection right after the duplication event and that when a new function is established, their evolution rate decelerates under purifying selection, losing the signatures of ancient positive selection due to the saturation of synonymous substitutions (Van de Peer et al. 2001; Jordan et al. 2004; Dong et al. 2012; Pegueroles et al. 2013).

The *OdsH* duplicate has been proposed to be a new gene in the *Sophophora* subgenus (Ting et al., 2004) and is associated with speciation in this clade, and we would not expect to see this gene further in *Drosophila* phylogeny. However, a search in the GenTree database (Shao et al. 2019) indicated the presence of *OdsH* in the ancestral node of the *Drosophila* genus, highlighting that its origin might be older than previously thought. We have thus asked the following questions: 1) how extensive is the presence of the *OdsH* duplicate in the phylogeny; 2) did neofunctionalization in testis occur before the divergence of the *melanogaster* subgroup; and 3) is *OdsH* associated with speciation in other recent species, beyond the *Sophophora* subgenus, such as the *mojavensis* complex (subgenus *Drosophila*)? We show that 1) the duplication occurred much earlier than previously proposed, dating back to 62 Mya in the Drosophilinae ancestor, 2) *OdsH* evolved under less intense negative selection than its paralogue *unc-4* and has features that allow the proposal of its ancient neofunctionalization in testis in the *Drosophila* genus, and 3) despite the presence and expression of *OdsH* in testis of the *mojavensis* complex, no clear association was established with the observed hybrid sterility in the crosses between species of this group.

## Results

### Occurrence and Phylogenetic Relationships

The search for sequences of the *unc-4* gene and its duplicate in annotated genomes resulted in sequences of 36 species of the genus *Drosophila* (subgenus *Sophophora* [25], *Drosophila* [9] and *Dorsilopha* [1]) and of *Scaptodrosophila lebanonensis* (Supplementary Table S1, Supplementary Material online). Only *D. melanogaster* and *D. sechellia* presented their predicted duplicates as *OdsH*. In the other species, the duplication was identified as *unc-4*, with the exception of *D. willistoni*, which is predicted to be a homologue of the gene homeobox pituitary Ptx1 (LOC6649333), and *D. albomicans*, which is predicted to be the segmentation polarity homeobox protein engrailed (LOC117569146). Sequences homologous to *unc-4* and duplicates were also obtained for groups of Drosophilinae underrepresented in the available annotated genomes (genus *Drosophila* [28] (subgenera *Sophophora* [10] and *Drosophila* [18]), *Zaprionus* [3], *Lordiphosa* [2], *Scaptomyza* [4] and *Chymomyza* [2]). With the exception of *D. erecta*, in whose genome the *OdsH* sequence was not found, the duplicate was found to be adjacent to the *unc-4* gene in the genomes of all species, and the synteny was conserved in this genomic region (encompassing *Socs16D*, CG12986 and *raskol* genes) (Fig. 1) along the *Drosophila* phylogeny. We also observed that the genomic fragment formed by the sequences of the genes *unc-4*, *OdsH* and *CG12896* probably underwent inversion in the *melanogaster* subgroup ancestor and in *D. takahashii* (Fig. 1). The investigated genomes from the subfamily Steganinae (4) returned only the *unc-4* sequence (Supplementary Table S2, Supplementary Material online). However, in *L. varia*, the only Steganinae representative that has genome assembled in contigs, the sequences of the reference neighbour genes (*Socs16D* and *raskol*) were found very far from the single copy *unc-4* sequence (*raskol* at 1.8 million base pairs and *Socs16D* at 4 million, both upstream), representing its neighbours *CG17209* upstream and *CG14213* downstream. No evidence of *unc-4* duplicates was found in genomes of the non-Drosophilidae Diptera (Supplementary Table S3, Supplementary Material online).

**Fig. 1.**
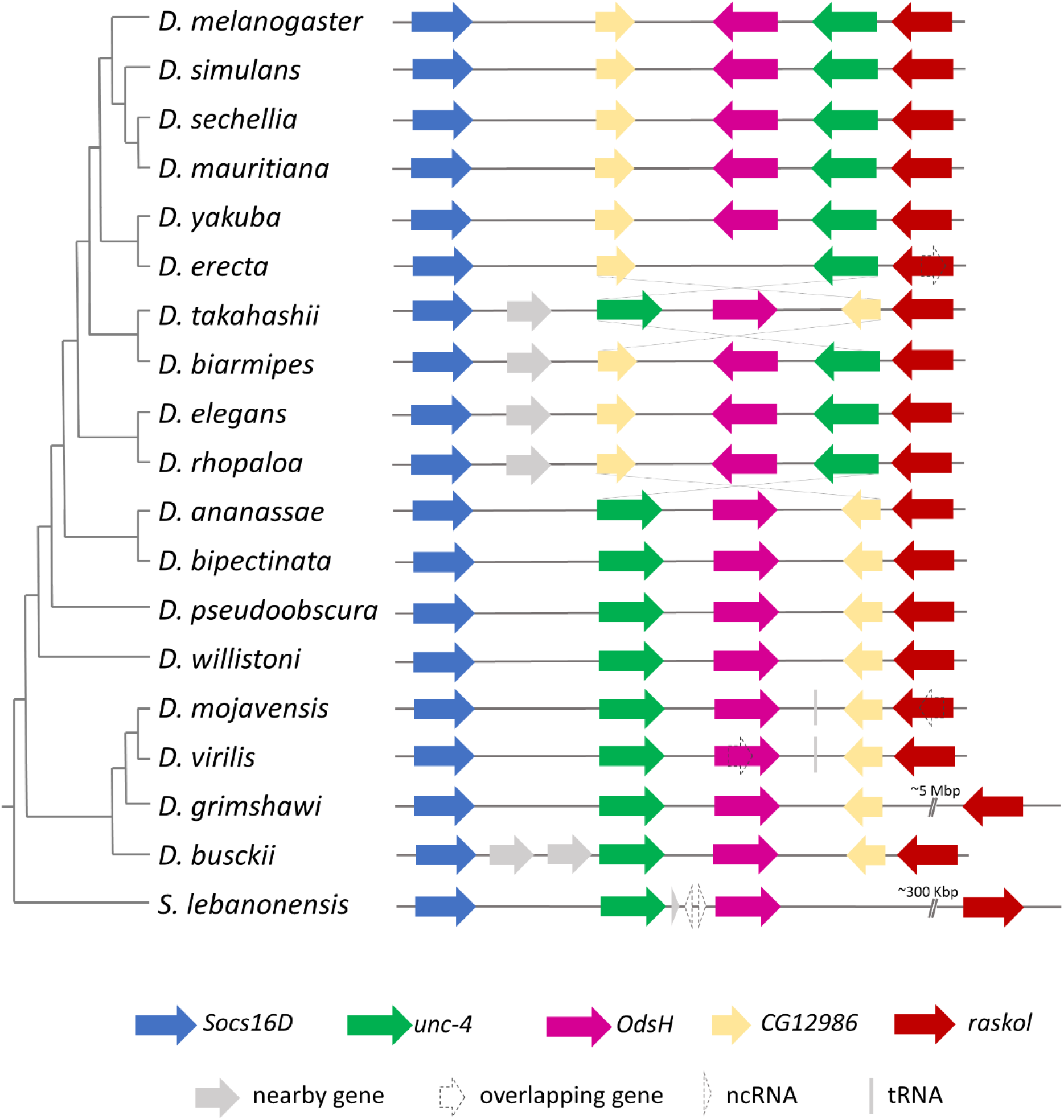
Relative positions of gene sequences in the neighbourhood of *OdsH* and *unc-4* in Drosophilinae genomes. The representation of the phylogenetic relationships is based on Suvorov al. 2022.

The distances between the duplicates varied between 10,982 bp (*D. simulans*) and 80,454 bp (*Scaptodrosophila lebanonensis*), and was 31,393 bp on average. The lengths of the *OdsH* genes ranged between 5,195 bp (*D. busckii*) and 37,364 bp (*D. willistoni*), with an average of 23,027 bp. The lengths of *unc-4* ranged between 7,801 bp (*D. willistoni*) and 21,691 bp (*D. virilis*), with an average of 11,536 bp. Although both genes present a general structure containing four exons (Fig. 2), they differ in size, mainly due to the longer introns in *OdsH*. Additionally, there is no signal of homology between their exon 1. Furthermore, *D. mojavensis*, *D. arizonae* and *S. lebanonensis* showed an extra exon upstream of the *OdsH* first exon, here referred to as exon 0. The same was observed for *unc-4* of *D. ananassae, D. virilis* and *D. grimshawi.* These extra exons probably arose independently in different evolutionary lineages since they show no homology among the orthologues from different groups (Fig. 2).

**Fig. 2.**
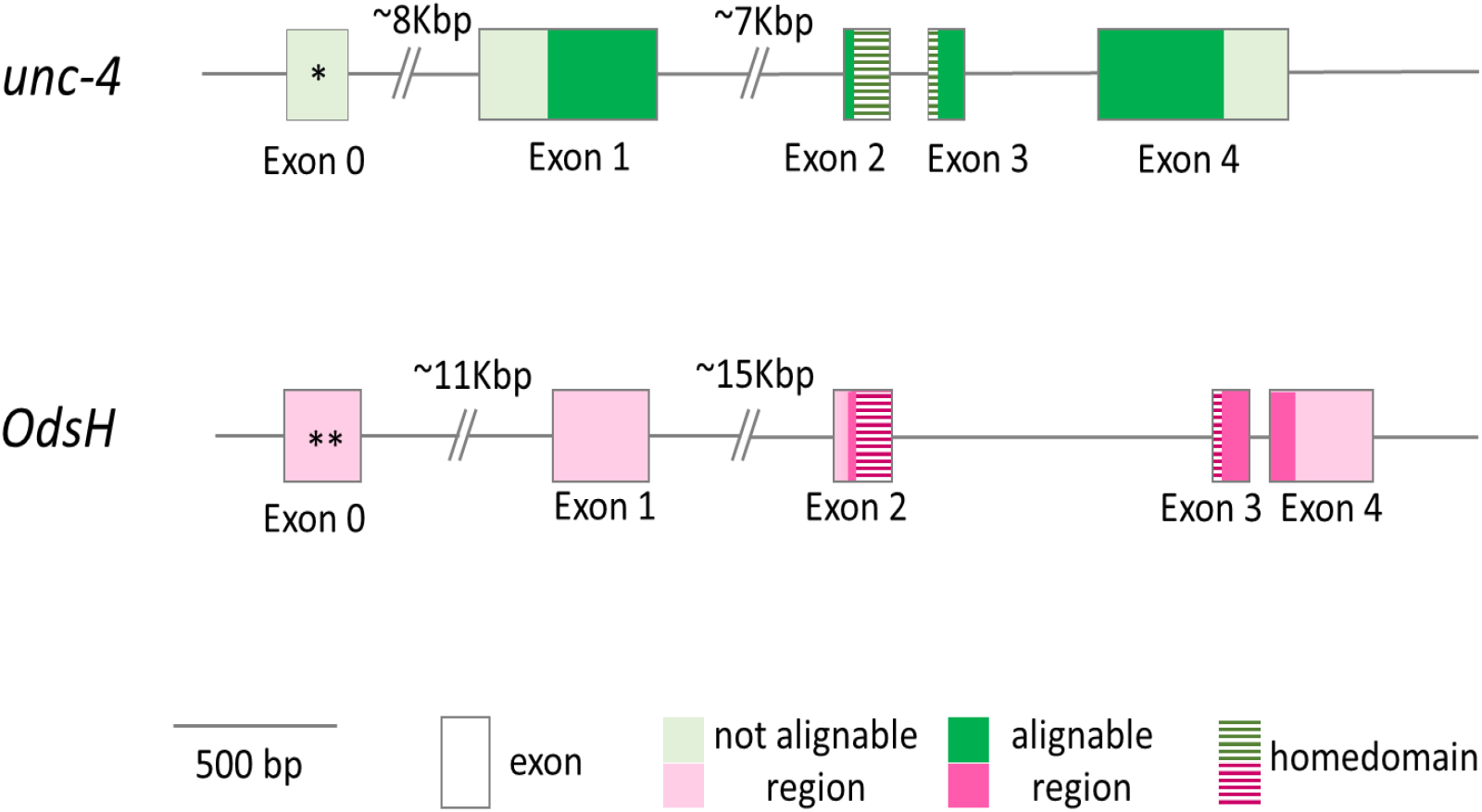
Gene structure of *unc-4* and *OdsH* in Drosophilinae. *Present only in *D. ananassae, D. virilis* and *D. grimshawi*. **Present only in *D. arizonae, D. mojavensis* and *S. lebanonensis*.

All *unc-4* and *OdsH* sequences used to infer the phylogenetic relationships between the two genes segregated into two sister monophyletic groups, supporting the hypothesis of orthology between the obtained *OdsH* sequences and the predicted *OdsH* of *D. melanogaster* and *D. sechellia*, as well as the paralogy in relation to *unc-4* (Supplementary Fig. S1, Supplementary Material online). Although the sequences of the *willistoni*-*saltans*-*Lordiphosa* radiation, which form a robust monophyletic cluster, coalesce to the common ancestral nodes in both the *unc-4* and *OdsH* clades, their positioning in both clades is inconsistent with the evolutionary history of Drosophilinae. This radiation grouped at the bottom of the Drosophilini branch for both genes. This incongruity may be due to the differential use of codons in this lineage in relation to the others, as already reported for the species of the groups *willistoni* and *saltans* (Rodríguez-Trelles et al. 2003; Vicario et al. 2007; Singh et al. 2006). We then calculated the *relative synonymous codon usage* (RSCU), the effective number of codons (ENC) and the percentage of GC content in the third position of the codons (%GC3). The Principal Component Analysis (PCA) of the RSCU data showed different codon usage patterns for *unc-4* and *OdsH* among species. For both genes, the *willistoni* and *saltans* groups, as well as the single copy *unc-4* of the Steganinae subfamily, were clustered with ∼40% variance from the *Drosophila* subgenus (Supplementary Fig. S2 and S3, Supplementary Material online). In addition, higher ENC values and lower %GC3 were observed in *unc-4* sequences from the *willistoni-saltans-Lordiphosa* branch in comparison to the other Drosophilini (ENC: *t* = -4.27, *p* = 3E-05; %GC3: *t* = 9.335, p < 0.00001) (Supplementary Fig. S5, Supplementary Material online) and in *OdsH* (ENC: *t* = -4.677, *p* = < 0.00001; %GC3: *t* = 9.884, p < 0.00001) (Supplementary Fig. S4, Supplementary Material online). Knowing that differences in the use of codons can cause phylogenetic artefacts (Inagaki et al. 2004, Inagaki and Roger, 2006; Liu et al. 2014), we removed these sequences from the phylogenetic analyses. In addition, sequences from groups of species that were clustered in the phylogeny in relation to the *Drosophila* subgenera incongruently to previous studies were also removed to avoid biases in the analyses of duplication dating and selection.

We used Bayesian inference to estimate the tree topology and the divergence time between *unc-4* and *OdsH* sequences of Drosophilinae. The monophyly of these genes was confirmed, building sister clades generally comprising the subgenera and species groups of Drosophilinae (Fig. 3, Supplementary Fig. S6, Supplementary Material online). The node shared by these two clades, which represents the duplication event, rooted by the *unc-4* single-copy sequences of the Steganinae clade, dated back to 62 million years ago. The *OdsH* clade has longer branches than *unc-4*, with older ages for the nodes of the taxa, an artefact due to the greater divergence between its sequences than between those of *unc-4*. However, the clades *OdsH* and *unc-4* show congruence regarding the monophyly of the tribes Drosophilini and Colocasiomyini and of the subgenus *Sophophora*, positioned basally in the tribe Drosophilini. The *unc-4* sequences of the genus *Zaprionus* were basally grouped with *Sophophora*, although without statistical support. Incongruences between the two branches were also observed regarding the internal organization of the clades of the subgenus *Drosophila* and other genera of Drosophilini, with *D. repletoides* grouped with *D. busckii*, positioned more basally than *Sophophora*, in the *unc-4* clade. However, inconsistencies were observed between the different phylogenetic proposals for the *Drosophila* subgenus (reviewed in O’Grady and DeSalle 2018).

**Fig. 3.**
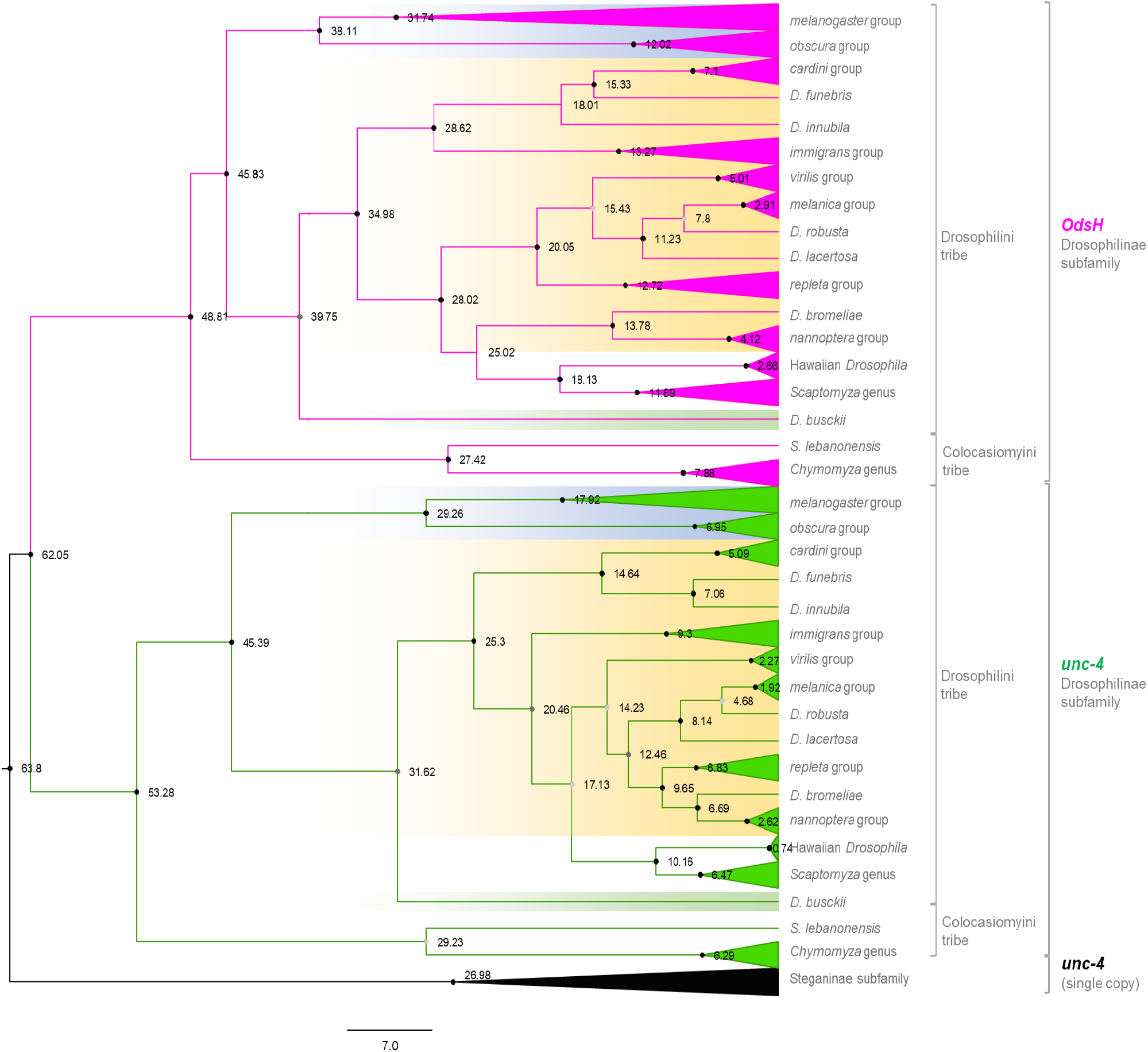
Calibrated Bayesian phylogenetic inference of the sequences of the paralogue genes *unc-4* and *OdsH* using the GTR+G+I substitution model. The analysis was performed with 405 nucleotide sites from 131 nucleotide sequences. All positions containing gaps and ambiguous bases were removed from the pairwise sequence analysis. The branches referring to the *Drosophila* taxonomic groups were compressed. At the root of each clade, the posterior probability is presented by black (>0.9) and grey (>0.7) circles, and the estimated times of divergence are indicated. The analysis was conducted in BEAST v16.1. The *unc-4* clade (green), subdivided into the more basal single-copy *Steganinae* (outgroup – black) and Drosophilinae, is presented at the base of the phylogeny followed by the *OdsH* clade in the upper part (pink). Monophyletic taxonomic groups of the *Drosophila* genus were compressed. Uncompressed clades can be seen in Supplementary Fig. S6, Supplementary Material online. Subgenera are highlighted in blue (*Sophophora*), yellow (*Drosophila*) and green (*Dorsilopha*). The brackets indicate tribes and subfamilies of Drosophilidae.

### Evolutionary Dynamics

The rate of nucleotide substitution was higher in *OdsH* than in *unc-4* (Z=8.395, p<0.05) in relation to the *unc-4* single copy of the outgroup. The signatures of selection on *OdsH* were estimated by the branch model – Model 2 (two ratio) by labelling each gene, using the tree estimated for them (*Tree 1*), and for each group of species represented by more than three sequences, with a tree estimated using only *OdsH* sequences (*Tree 2*) and for specific branches (*D. simulans*, *D. sechellia* and *D. mauritiana*, and *D. navojoa* and *D. mojavensis*) that presented ω>0.3 in the branch Model 1 analysis. No nonsynonymous substitutions were detected in any *unc-4* sequence in the Drosophilidae family, and therefore analysis of selection was not performed for this gene. Negative selection was predominantly observed in the evolution of the two genes (ω<1) in the branch Model 2 analysis; however, the mean values of ω differed significantly (χ² = 10.266, p = 0.001), being approximately four times higher for *OdsH* (average ω = 0.196) than for *unc-4* (ω = 0.052) and many times greater if we consider the value of ω only in Drosophilinae, which is zero, as there are no nonsynonymous substitutions between the sequences of *unc-4* in this subfamily. A single nonsynonymous substitution was observed between these sequences and that of the outgroup *Rhinoleucophenga bivisualis* (T118Q). The labels in the *OdsH* ancestral node and along its branches showed higher ω values than the background, ω = 0.204 (χ² = 10.092, p = 0.001) and ω = 0.049 (χ² = 113.817, p = 0), respectively, indicating greater divergence in relation to *unc-4* in the ancestral node *OdsH* than along its divergence. As no nonsynonymous substitution was observed in the Drosophilinae *unc-4* sequences, branch-model analysis was not performed for this gene. For the selection acting on *OdsH*, no differences were observed between the groups of Drosophilinae species (Table 1), except for sequences of the Hawaiian *Drosophila* (ω = 0.173; χ² = 4.044, p = 0.044) and the *D. melanogaster* group (ω = 0.023, χ² = 4.25, p = 0.039). Although the *D. melanogaster* group has a ω value lower than that of the sequences from the other groups, the intensity of negative selection is relaxed for the clade *D. simulans*–*D. sechellia*–*D. mauritiana* (ω = 0.611; χ² = 30.84, p = 0). In addition to these species, labels were specified in the branches of *D. navojoa* (ω = 0.248) and *D. mojavensis* (ω = 0.458), whose differences were not significant in relation to the background (respectively χ² = 1.031, p = 0.31, and χ² = 3.352, p = 0.067), although *D. mojavensis* showed a ω indicative of relaxed selection. No signals of sites under positive selection in *OdsH* were detected (Supplementary Table S4, Supplementary Material online).

**Table 1.** Selective process acting on *OdsH* in groups and species of Drosophilinae, estimated through the branch model (*two ratio*).

| Groups of species | $\omega_0$ | $\omega_1$ | lnL | np | $\chi^2$ | df | P |
| --- | --- | --- | --- | --- | --- | --- | --- |
| M0 | 0.051 |  | -5315.46 | 118 |  |  |  |
| ***Melanogaster*** | **0.053** | **0.023** | **-5313.336** | **119** | **4.250** | **1** | **0.039** |
| ***D. simulans* -**<br>***D. mauritiana* -**<br>***D. sechellia*** | **0.047** | **0.611** | **-5300.040** | **119** | **30.843** | **1** | **0** |
| *obscura* | 0.051 | 0.052 | -5315.461 | 119 | 0.001 | 1 | 1.000 |
| *repleta* | 0.051 | 0.053 | -5315.461 | 119 | 0.001 | 1 | 1.000 |
| *D. navojoa* | 0.051 | 0.248 | -5314.946 | 119 | 1.031 | 1 | 0.310 |
| *D. mojavensis* | 0.051 | 0.458 | -5313.785 | 119 | 3.351 | 1 | 0.067 |
| *virilis* | 0.052 | 0.019 | -5314.932 | 119 | 1.058 | 1 | 0.304 |
| **Hawaiian *Drosophila*** | **0.050** | **0.173** | **-5313.439** | **119** | **4.044** | **1** | **0.044** |
| *Scaptomyza* | 0.052 | 0.018 | -5314.818 | 119 | 1.285 | 1 | 0.257 |
| *immigrans* | 0.052 | 0.040 | -5315.406 | 119 | 0.111 | 1 | 0.740 |
| *Chymomyza-Scaptodros*<br>*ophila* | 0.052 | 0.023 | -5314.809 | 119 | 1.305 | 1 | 0.254 |

M0: null hypothesis, without phylogenetic labelling; $\omega_0$ : Ka/Ks ratio of the group outside the label; $\omega_1$ : Ka/Ks ratio of the labelled group; np: number of parameters; df: degrees of freedom. Groups that have different $\omega$ values are highlighted in bold.

### Candidate Regulators of *OdsH* and *unc-4* Expression

The comparison of the 500 bp upstream and downstream regions of *OdsH* and *unc-4* showed that *OdsH* was enriched for 43 and 15 transcription factor-binding sites (TFBSs), respectively, while *unc-4* upstream and downstream regions had 15 and 13 TFBSs, respectively (Fig. 4; Supplementary Tables S5 and S6, Supplementary Material online). Transcription factors that putatively bind to the regulatory region of *OdsH* showed a wide diversity of GO categories primarily related to development and organogenesis, while those of *unc-4* were also related to leg development and morphogenesis (Supplementary Fig. S7, Supplementary Material online). In the upstream region of *OdsH*, the enrichment of TBFSs attributed to the category of development process involved in male reproduction stood out, specifically *achi*, *vis* and *so*, which are related to the spermatogenesis category.

**Fig. 4.**
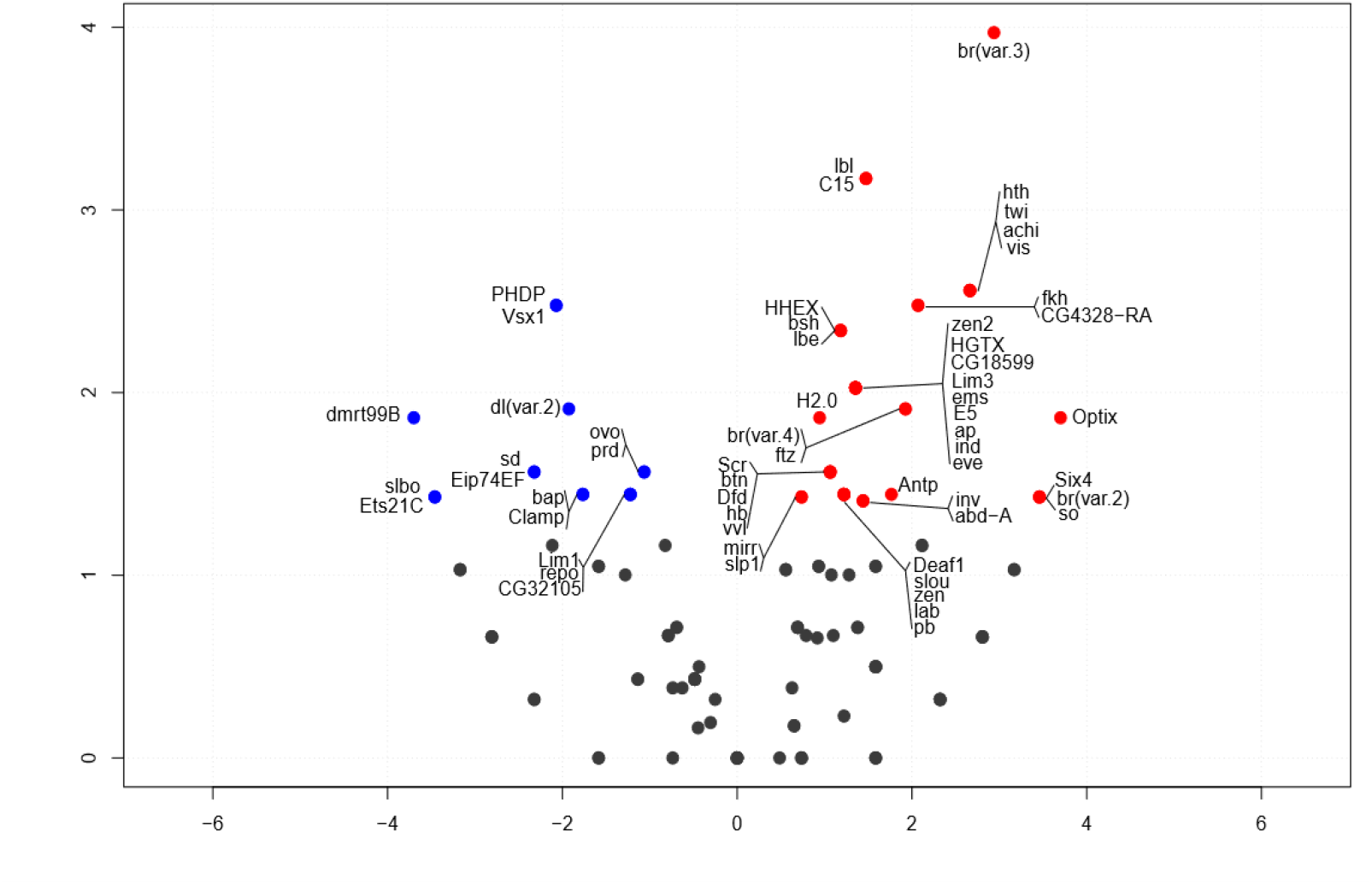
Enrichment of transcription factors that had binding sites enriched for *OdsH* are represented in red and for *unc-4* are represented in blue. Grey dots represent transcription factors whose binding sites did not differ from each other.

### Functional Protein Motifs

The homeodomain and the C-terminal octapeptide were conserved in the sequences of the Unc-4 proteins (e-values – homeodomain: 1.0e-3418; octapeptide: 5.8e-238) and OdsH (e-values – homeodomain: 2.3e-3505; octapeptide: octapeptide: 1.6e^-189^) in *Drosophila*, as seen in the scheme of primary structures in *D. melanogaster* (Fig. 5). In both motifs, there was greater divergence in OdsH, while the Unc-4 motifs did not show amino acid substitutions (Fig. 5 A). The OdsH octapeptide has a core of eight conserved amino acids, and the adjacent amino acids exhibit some divergence. OdsH in *D. mauritiana* is missing the octapeptide, since there is a truncation at the C-terminal region. The three-dimensional models of the homeodomains showed the usual secondary structure of three alpha helices with an N-terminal tail in a segment of 54 amino acid residues (Fig. 5 B), with the exception of amino acid 53 at the C-terminal end of the third helix in OdsH. In Unc-4, this amino acid does not participate in the structure. Since the Unc-4 homeodomain did not have substitutions in *Drosophila* or in *T. dalmani*, there was no variation in the free energy of protein/DNA binding (ΔΔGb). Conversely, OdsH homeodomains showed higher DNA binding instability, which was more pronounced in *D. simulans* (−7.896 kJ/mol) and *D. mauritiana* (−7,414 kJ/mol) (Fig. 5 C). Most OdsH homeodomain substitutions destabilized the complex with DNA (ΔΔGb<0) (Fig. 5 D). It was generally observed that the species had different substitutions in OdsH that resulted in different ΔΔGb per site, except for *D. persimilis* and *D. pseudoobscura*, which have identical sequences, and *D. mojavensis*, *D. virilis* and *D. grimshawi*, which have similar numbers of amino acid substitutions (6 substitutions in *D. mojavensis* and *D. virilis*, and 7 in *D. grimshawi*, 4 of which were shared between the three species). The species in the *melanogaster* subgroup had substitutions that resulted in the highest ΔΔGb values. A greater number of substitutions was found in the first α-helix. In the third α-helix, which makes direct contact with the DNA, there were two substitutions shared by different groups (S40G, except for *D. simulans*, *D. mauritiana* and *D. sechellia*, which shared the ancestral allele, and V53W). The other substitutions in this helix were species specific and were present exclusively in the *melanogaster* group.

**Fig. 5.**
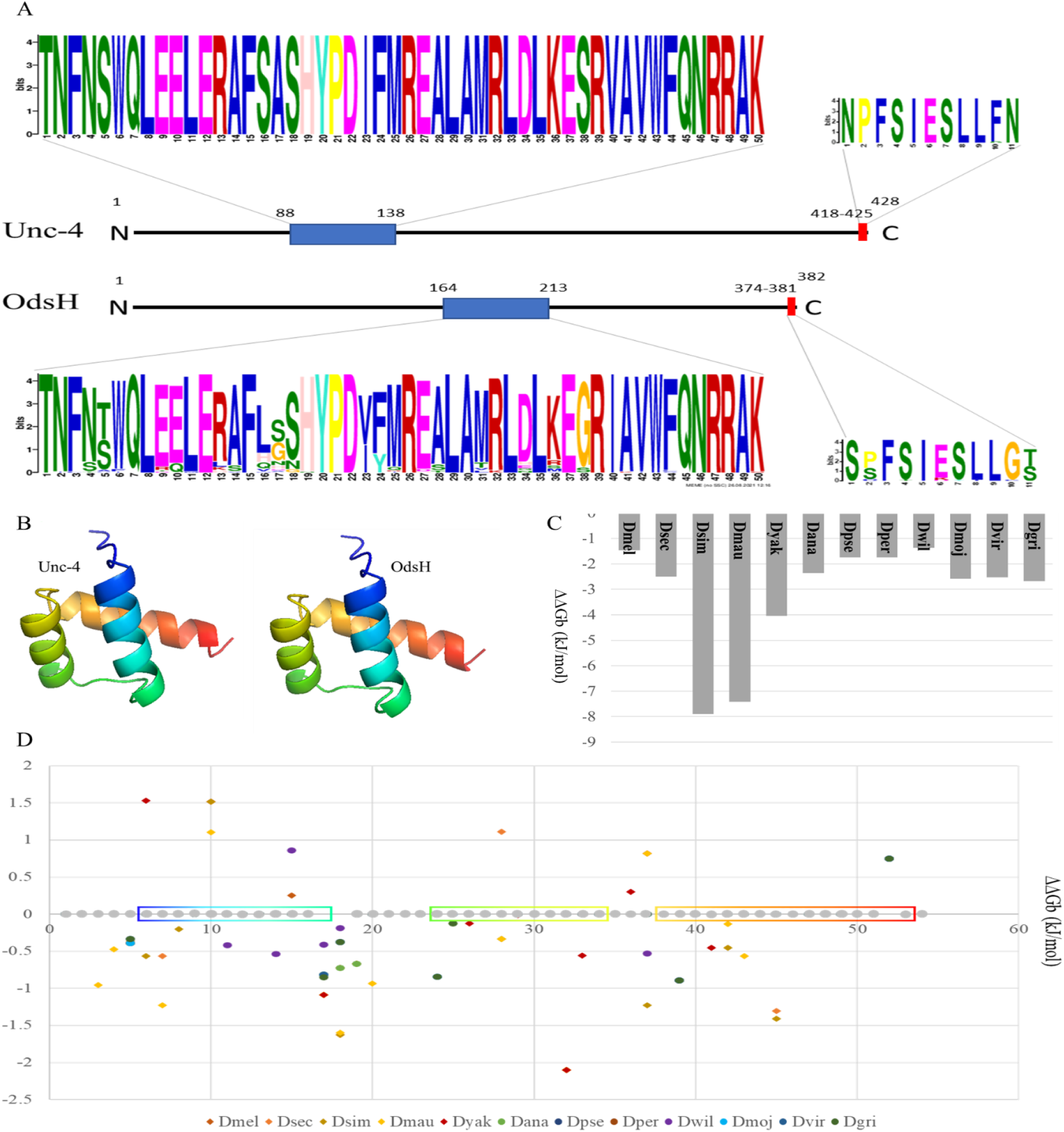
Functional motifs in Unc-4 and OdsH proteins. A. Representations of the Unc-4 and OdsH primary structures in *D. melanogaster* and functional motifs found in Drosophilinae: homeodomain (blue) and octapeptide (red). B. Three-dimensional models of Unc-4 and OdsH homeodomains. The N-terminal tail is presented in blue, and the C-terminal tail is presented in red. C. Total energy variation of the OdsH and DNA homeodomain complex, by species, in relation to Unc-4. D. Energy variation of the OdsH and DNA homeodomain complex, per substitution, relative to Unc-4, by species along the amino acid chain (0-54). Sites without a grey circle represent replacement in all analysed species. The boxes represent the positions of the 3 α-helices. Overlapping dots represent shared mutations: 5-*D. mojavensis*, *D. virilis*, *D. grimshawi*; 5-*D. persimilis*, *D. pseudoobscura*; 17-*melanogaste*r complex; 17-*D. persimilis*, *D. pseudoobscura*, *D. ananassae*, *D. mojavensis*, *D. virilis*; 18-*D. sechellia*, *D. simulans*; 18-*D. persimilis*, *D. pseudoobscura*, *D. mojavensis*, *D. virilis* and *D. grimshawi*; 19-*D. simulans*, *D. mauritiana*, *D. yakuba*, *D. ananassae*; 32-*melanogaster* complex; 37-*D. melanogaster*, *D. sechellia*, *D. mauritiana*; 39-all except *D. simulans*, *D. sechellia* and *D. mauritiana*; 52-all species.

### Expression of *OdsH* and *unc-4* in *D. arizonae*, *D. m. baja* and their Hybrids

The analysis of the *Drosophila* transcriptomes available in public databases *(D. pseudoobscura*: PRJNA291085; *D. grimshawi*: PRJNA317989: *T. dalmani*: PRJNA240197; other species: PRJNA388952) showed that both genes have low expression levels. However, *unc-4* seems to be mainly expressed in somatic tissues, whereas *OdsH* seems to be specific to male reproductive tissues (Supplementary Fig. S8, Supplementary Material online). This is expected in the cases of neofunctionalization, suggesting that *OdsH* neofunctionalization occurred rapidly after its origin.

To identify whether the expression of *OdsH* in the testis of sterile hybrids is atypical in other *Drosophila* groups, as described for the crosses between *D. mauritania* and *D. simulans*, we analysed species from the *repleta* group that show incipient speciation. We performed single-molecule RNA fluorescence *in situ* hybridization (smRNA FISH) of *OdsH* in the testes of *D. arizonae* and *D. mojavensis baja* species and their respective hybrids, since their hybrids present a sterile or fertile phenotype depending on the cross direction. During spermatogenesis, spermatocytes are known to show an increase in cell and nuclear volume and open chromatin (Fig. 6 A). We observed *OdsH* transcripts in the primary and secondary spermatocytes in the parental strains (Fig. 6 B-E). The patterns of the spermatocyte staining do not seem to be different from the parental ones in both H♀moj^baja^♂ari (fertile) (Fig. 6 F) and H♀ari♂moj^baja^ (sterile) (Fig. 6 G) hybrids. In addition, no signal of *OdsH* expression was observed in cells at the extreme apex of the testes or in the postmeiotic stages. Furthermore, we could observe that the sterile hybrids differ from the fertile ones by the defective formation of the sperm bundles (Supplementary Fig. S9, Supplementary Material online).

**Fig. 6.**
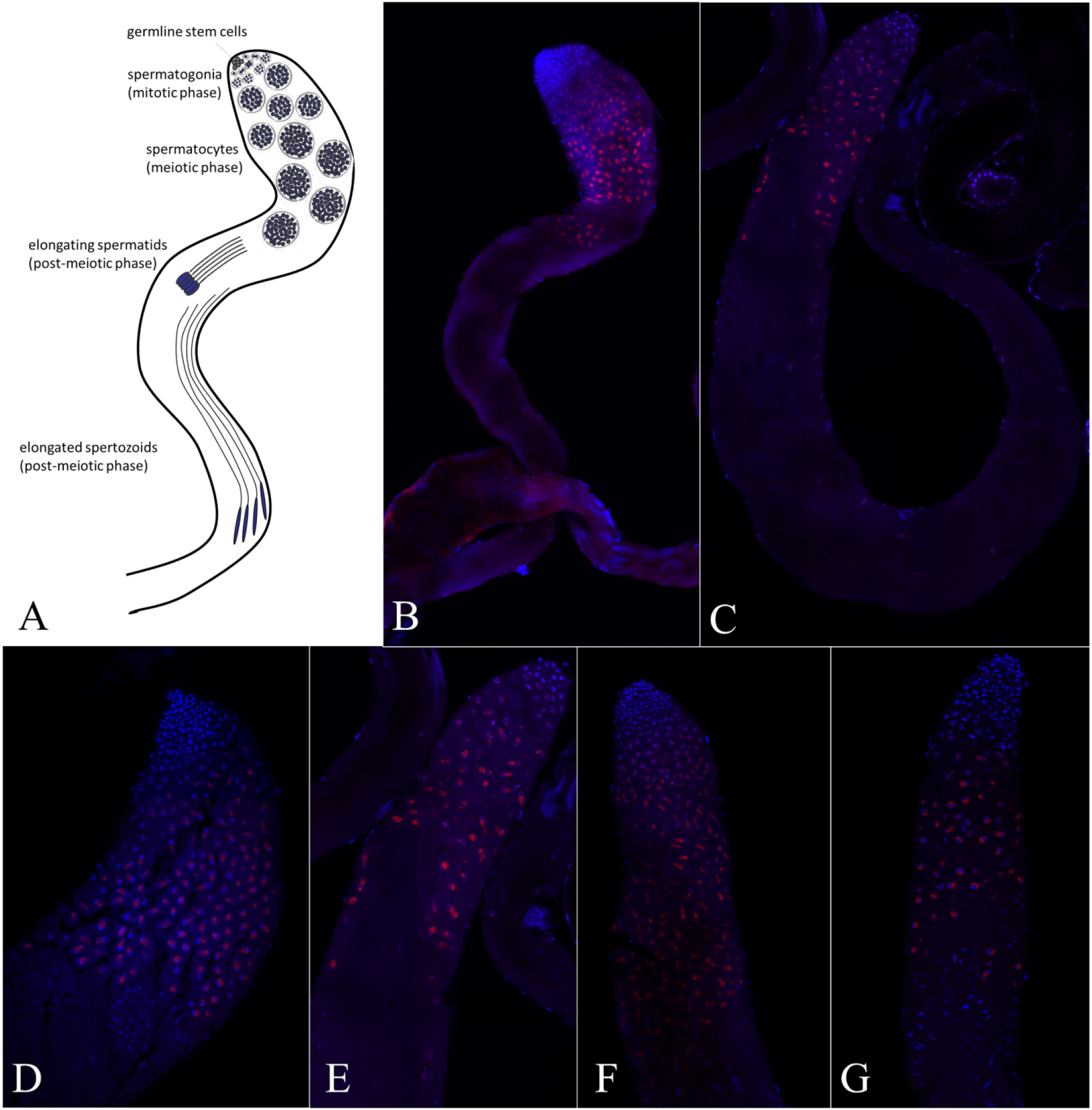
Single-molecule RNA fluorescence *in situ* hybridization (smRNA FISH) of *OdsH* in testes of *D. arizonae*, *D. m. baja* and its hybrids. A. Scheme of *Drosophila* spermatogenesis. B. Panorama of testes of *D. m. baja*. B. Panorama of testes of *D. arizonae*. C. Apical region of testis of *D. m. baja*. D. Apical region of testis of *D. m. baja*; E. Apical region of H♀moj^baja^♂ari (fertile) testis. F. Apical region of H♀ari♂moj^baja^ (sterile) testis. Notes – blue: DAPI; red: *OdsH* probes. H: hybrid.

To identify whether the expression of *OdsH* could be quantitatively differentiated in these hybrids, we quantified its expression in the testes of the parental species and their hybrids by RT-qPCR (Supplementary Table S7, Supplementary Material online). The levels of expression were different (KW= 22.24, p<0.001) between parental species but not between hybrids or between parental strains and hybrids, except for the comparison of *D. m. baja* and H♀ari♂moj^baja^ (Supplementary Fig. S10, Supplementary Material online).

## Discussion

### The Emergence of a New Duplicate in the Drosophilinae Subfamily

The hypothesis of the *OdsH* origin from a duplication of the *unc-4* gene in the *Sophophora* subgenus ancestor was proposed by Ting et al. (2004). It was based on the presence of this gene in species of the *melanogaster* and *obscura* groups (*Sophophora* subgenus) without dating the duplication. To answer this question, we looked for sequences homologous to *unc-4* in all available genomes of the Drosophilidae (Bächli 2016). We identified *unc-4* duplicates in genomes from six genera of the Drosophilinae subfamily (*Drosophila*, *Scaptodrosophila*, *Chymomyza*, *Scaptomyza*, *Lordiphosa* and *Zaprionus*) but not in Steganinae or other families of Diptera. This suggests that the duplication occurred much earlier than previously suggested by Ting et al. (2004) and placed the duplication in the ancestor of the subfamily Drosophilinae. We cannot exclude the possibility that the duplication occurred in a more basal node and was lost in other branches, but we do not have any argument to support this scenario.

No evidence of *unc-4* duplicates was observed in the genome of *D. erecta*, which may be due to artefacts in the assembly of the genome (Denton et al. 2014). However, we cannot rule out the possibility of loss in this species. Loss of one of the copies due to accumulation of random mutations is a common fate among duplicated genes (Ohno 1970; Wolfe and Shields 1997; Inoue et al. 2015). Moreover, in *D. melanogaster*, the knockdown of this duplicate has no effect on the individual’s viability (Sun et al. 2004; Cheng et al. 2012).

Because the orthology of each duplicate and the paralogy between them are supported by the *in tandem* positioning in the assembled genomes (Fig. 1) and the phylogenetic relationships (Fig. 3), which are robust evidence of homology (Altenhoff and Dessimoz 2012), we considered that the duplicated gene is *OdsH*. By using a Bayesian phylogenetic inference approach, we conclude that *OdsH* and *unc-4* belong to sister monophyletic clades, which is evidence of a unique evolutionary origin of *OdsH* in Drosophilinae.

The presence of *OdsH* exclusively in Drosophilinae and in all its subgenera indicates that duplication occurred in the ancestral lineage of this subfamily at an estimated time of 62 Mya, right after the spread of the ancestor lineages of the subfamilies Steganinae/Drosophilinae. Suvorov et al. (2022), using genomic data, developed a broad dating analysis of Drosophilidae, whose divergences were estimated to be 63.2 Mya. The estimate for the divergence of the Drosophilinae subfamily in our analysis (53.3 Mya in the *unc-4* clade and 48.81 Mya in the *OdsH* clade) is close to that proposed by Suvorov et al. (2022) (53.4 Mya).

### *OdsH* and *unc-4*: Same Origin, but Divergent Evolutionary Histories

The sequences of *OdsH* and *unc-4* have evolved asymmetrically, since the former shows a higher divergence along Drosophilinae. *OdsH* shows more indels and thus smaller regions that can be aligned between the orthologous sequences in comparison to *unc-4* (Fig. 2).

*OdsH* showed higher rates of amino acid replacements and less intense negative selection on the Drosophilinae ancestor node than its paralogue *unc-4*. This indicates faster evolution of *OdsH* in the Drosophilinae ancestor. Along its divergence, we estimate stronger negative selection, in agreement with the scenario of ancient neofunctionalization proposed by Van de Peer et al. (2001). The signatures of negative selection were homogeneous in the Drosophilinae, except in the Hawaiian *Drosophila* and *melanogaster* group (Table 1), which showed relaxation of negative selection on the branch formed by *D. simulans-D. mauritiana-D. sechellia*. However, our branch model analyses did not identify positive selection as in the pairwise model reported by Ting et al. (1998).

In addition to sequence and phylogenetic divergence, we did not observe the presence of *unc-4* expression in the gonads of males (except for *D. yakuba* and *D. ananassae*) and females. *unc-4* is conserved in Metazoa and is related to the analysed species, in agreement with the data observed for the single copy of the outgroup *Teleopsis dalmani*. This functional conservation is also supported by its lower diversity of putative TFBSs (Fig. 4) and lack of amino acid replacements in its homeodomains and octapeptides in Drosophilinae when compared to the single-copy gene in Steganinae, indicating energy stability of homeodomain binding to DNA (Fig. 5).

Regarding *OdsH*, by using public datasets from NCBI, we observed expression exclusively in male reproductive tracts and testes in *Drosophila*, except for *D. pseudoobscura* (Supplementary Fig. S8, Supplementary Material online). We also found that *OdsH* expression levels were higher (from 169.5 to 340 normalized read counts) than *unc-4* expression levels (less than 50 normalized read counts) (Banho et al. 2020) in transcriptomes of the reproductive tracts from two *D. mojavensis* subspecies and *D. arizonae* previously sequenced by our group (BioProject NCBI PRJNA691040). Additionally, the expression levels of both genes in the female reproductive tract were lower than 10 counts (Banho et al. 2021).

In contrast to *unc-4*, the *OdsH* sequence was enriched in a greater diversity of TFBSs in its regulatory regions (Fig. 4), which is in agreement with the observation of higher complexity in the regulatory regions of ancient daughter duplicates during their divergence (Zhang and Zhou 2019). In addition, TFBSs related to the development of the male reproductive system and to the initial stages of spermatogenesis (*achi*, *so* and *vis*) were enriched in *OdsH*. It is known that *achi* and *vis* are expressed in primary spermatocytes, acting on the specification of the spermatogenesis gene regulation program (Ayyar et al., 2003; Wang and Mann 2003). Moreover, it has been shown that *so* is expressed in the cyst cells of the apical region of the *Drosophila* testis and contributes to the normal development of primary spermatocytes (Fabrizion et al. 2003).

Particularly with respect to sequence divergence, the OdsH protein shows greater divergence of the homeodomain than Unc-4, which can disturb the DNA binding energy, making the system more unstable (Fig. 5). These particularities of OdsH might make the binding of its homeodomain to its DNA target sites less specific than that of Unc-4. This suggests that the two proteins, which are transcription factors, have different binding sites in the target DNA that they regulate. However, OdsH, like Unc-4, has the conserved homeodomain amino acid Q47, which gives high cooperativity to homeodomains, with cooperativity being the main factor involved in the specificity of homeodomain binding to DNA target sites (Wilson 1995). The amino acids that directly interact with the nitrogenous bases of DNA are also conserved in OdsH and Unc-4 (V44 and N48 [Wilson 1995]), with the exception of *D. mauritiana*, which has an isoleucine at residue 44 of the OdsH homeodomain.

In view of the evolutionary changes discussed above, we propose that neofunctionalization of *OdsH* occurred in the testes of the Drosophilinae ancestor. Subsequently, *OdsH* seems to have evolved different functions in the Drosophilinae evolutionary lineages, since it seems to be expressed not in testes but in other reproductive and nonreproductive tissues in the *obscura* (*D. pseudoobscura* and *D. persimilis*) and *virilis* (*D. virilis*) groups of *Drosophila*. Our observations are in agreement with the out-of-testes hypothesis (Kaessman 2010) and with previous reports of the acquisition of new functions of newly duplicated genes in *Drosophila* testes (Assis and Bachtrog 2013; Assis 2014), with posterior evolution of out-of-testes functions along the divergence time (Kaessman 2010), accompanied by higher sequence conservation (Jiang and Assis 2017). The dating of the duplication that originated *OdsH* at 62 Mya ago and our hypothesis of early neofunctionalization finds support in Bao et al. (2018), who demonstrated that duplicates in *Drosophila*, dated to approximately 60 million years ago, underwent higher rates of neofunctionalization and innovative evolutionary rates. This may have configured a propitious scenario for fixing substitutions and neofunctionalization at the time of *OdsH*/*unc-4* duplication.

### The role of *OdsH* as a Speciation Gene

Our protein sequence analysis identified the replacement of the amino acid valine, conserved at site 44, which interacts directly with nitrogenous bases of its binding site in DNA, by isoleucine in the *OdsH* of *D. mauritiana* (Fig. 5 C). This might change the binding sites on the genome. It was previously identified that *D. mauritiana OdsH* binds to the heterochromatic region of the Y chromosome, whereas that of *D. simulans* does not bind to this region (Bayes and Malik 2009), which may be caused by this difference in the DNA strand-binding amino acid in *D. mauritiana*. Moreover, the OdsH proteins in the two species are the ones with the highest values of DNA-binding instability (Fig. 5 C), probably driven by the relaxation of negative selection observed in these sequences.

Additionally, the specificity of binding to sites on the DNA strand depends mainly on transcription cofactors that act linked to homeodomains (Wilson et al. 1995; Bürglin and Affolter 2016). The evolution of homeodomains of the paired-like phylogenetic class, to which Unc-4 (Winnier et al. 1999) and *OdsH* belong, occurs through rearrangements and losses of functional motifs, including the octapeptide. The diversity of protein structures in this class of proteins is mainly related to the presence/absence of functional motifs between its families (Jacob 1977). Indeed, the presence of the octapeptide is conserved in Unc-4 of *C. elegans* and binds to the transcription cofactor Unc-37 (orthologous to Groucho, in *Drosophila*), repressing its target expression (Winnier et al. 1999).

Our analyses also showed that OdsH of *D. mauritiana* does not show the octapeptide, which is conserved at the C-terminal ends of Unc-4 and OdsH of the other Drosophilinae. Since the *OdsH* molecular mechanism of action occurs through the interaction of different loci (Bayes and Malik 2009; Lu et al. 2010), the structural features of the OdsH protein from *D. mauritiana* might result in incompatibility within the *D. simulans* genome, as proposed by the Dobzhansky-Muller model (Dobzhansky 1937; Muller 1942). This incompatibility leads to the phenotype of defective sperm bundle formation, resulting in immobility (Lu et al. 2010).

We previously observed sperm immobility in sterile hybrids of *D. arizonae* – *D. mojavensis* (Banho et al. 2021), and defects in sperm bundles have also been observed (Supplementary Fig. S9, Supplementary Material online; Kanippayoor et al. 2020). In these species, we showed that *OdsH* expression occurs during the differentiation of spermatocytes (Fig. 7), in which intensive cell growth and greater synthetic RNA activity occur (Hackstein 1987). In addition, for these species, the nucleus of mature primary spermatocytes has been described as dumbbell-shaped (Pantazidis et al. 1992), in which we can observe the highest intensity of *OdsH* probes (Fig. 7). Thus, our results indicate that *OdsH* is expressed in spermatocytes, as previously demonstrated in *D. simulans*, and its expression occurs in spermatocytes beginning in the G2 phase (Bayes and Malik, 2009). However, this feature is observed in a reduced number of old gene duplicates, such as *OdsH*, which are mostly expressed in the mitotic phases of spermatogenesis (Raices et al. 2019; Su et al. 2021).

In contrast to the atypical intense expression of *OdsH* in the apical cells of the testes in the sterile offspring from *D. mauritiana* and *D. simulans* (Sun et al. 2004), we showed that the expression of *OdsH* in *D. arizonae*, *D. m. baja* and their sterile and fertile hybrids did not differ (Fig. 7 and Supplementary Fig. S10, Supplementary Material online). This indicates that *OdsH* probably does not act as a speciation gene in these sister species. Indeed, speciation genes have been characterized as lineage specific (Gomes and Civetta, 2014), and *OdsH* might act as a speciation gene only in *D. mauritiana* and *D. simulans*.

In conclusion, we show here an older origin of *OdsH* than previously reported and the evolutionary process this duplicate underwent in Drosophilinae, as it evolved asymmetrically in relation to its ancestor gene *unc-4* and accumulated changes that resulted in neofunctionalization. We also report specific features that indicate protein divergence, particularly in *D. mauritiana*, which may be associated with the incompatibility described in introgression of this gene in the *D. simulans* genomic background. Our data show that even though it is the first speciation gene described in *Drosophila*, much of the evolutionary history that led *OdsH* to play a role in reproduction remains unknown and that its role as a speciation gene may be restricted to specific groups of species. The extent of such a role in the family Drosophilinae can only be determined with extensive studies using interspecific hybrids of closely related species similar to ours.

## Materials and Methods

### *unc-4* and *OdsH* annotation in the Drosophilidae genomes

The sequences of *unc-4* and its duplicates were retrieved from publicly available annotated Drosophilidae genomes, focusing on its two sister subfamilies, Steganinae and Drosophilinae, with BLAST (NCBI), selecting the *High Scoring Pairs* (HSPs) (Supplementary Table S1, Supplementary Material online). The mRNA sequences with the highest *scores* and *e-values* smaller than 1-e05 were aligned with MAFFT (Katoh et al. 2002). The alignments were verified with BioEdit Sequence Alignment Editor v. 7.0.9 (Hall 1999) to remove the sequences that did not align and nonhomologous regions with *indels*. Therefore, the aligned sequences included a conserved region among the duplicates that contains the homeodomain (162 bp), with 15 bp upstream of the N-terminal homeodomain end and 228 bp downstream of the C-terminal end. The conserved region found in the *D. melanogaster* duplicates was used as a *query* with BLAST to search for homologous regions with genomes that did not have their respective annotations (Supplementary Table S2, Supplementary Material online) using the same parameters as described for annotated genomes with a script in BASH language written by our group. Sequences from *Chymomyza procmenis, Cacoxenus indagator* and *Rhinoleucophenga bivisualis* were annotated in their genomes and assembled using SPADES v.3.9.0 software (Bankevich et al. 2012). For the annotation, the amino acid sequences of OdsH and Unc-4 of *D. melanogaster* were used as queries in TBLASTN searches in assembled genomes, and the scaffolds containing both homologous gene sequences were investigated on the coding sequences using the software Genewise (Birney et al. 2004). Analysis of synteny was performed manually considering the Drosophilidae genomes available in the OrthoDB database (Zdobnov et al. 2021). For *S. lebanonensis* and *L. varia*, which are not available in OrthoDB, BLASTX (Altschul et al. 1997) was used on the *D. melanogaster* protein database, considering a threshold of 70% or similarity and coverage. In addition, we also looked for *unc-4* and possible duplicates in the publicly available genomes of Diptera, which are outgroups of Drosophilidae (Supplementary Table 1, Supplementary material online). Since we found only a single copy duplication in these taxa, as in Steganinae, we decided to use only Steganinae data as the duplication outgroup for further analysis.

### Phylogenetic inference and duplication dating

Duplication dating was carried out with the Bayesian molecular clock method and lognormal transformation to estimate the consensus tree topology and the divergence time. It was possible to set the monophyly between *unc-4* and *OdsH* in Drosophilinae, as there is no evidence supporting duplicates *in tandem* in external taxa from such divergence. This method was used to avoid the phylogenetic bias *long branch attraction* (LBA) (Felsestein 1978; Hendy and Penny 1989), which has been demonstrated previously in phylogenetic heuristic methods with paralogues that have asymmetric evolution in *Drosophila* (Bao and Friedrich, 2009). Therefore, this method was used under the hypothesis that *unc-4* and *OdsH* evolved at different rates in comparison to the single-copy *unc-4* outgroup in Steganinae. Therefore, it could cause branch attraction in the most conserved gene, repulsion to the clade with the most divergent duplicate (LBA) and artefacts in the estimated dates.

Conserved region alignment was used to perform Bayesian inference of the phylogenetic relationships by the Yule process (Yule 1925; Gernhard 2008). For this, the software BEAST v. 1.6.1 (Drummond et al. 2006) was used with five categories of gamma distribution, invariable sites and the substitution model GTR (Nei and Kumar 2000), estimated as the best substitution model by BIC on MEGA X (Kumar et al. 2018). The dating was carried out using the lognormal relaxed molecular clock (Drummond et al. 2006). The calibration was assessed using the estimated divergence from Suvorov et al. (2022) as the calibration points, as their report presents intermediate ages for Drosophilidae branches in comparison to previous studies: Drosophilidae family ancestor (63.19 Mya, CI 95%: 58.79-65.73 Mya), Drosophilini tribe ancestor (46.84 Mya, CI 95%: 43.85-49.85 Mya) and *D. melanogaster* x *D. simulans* divergence (3.62 Mya, CI 95%: 2.92-4.40 Mya) in the divergence node of its respective groups at the *unc-4* and *OdsH* clades. This calibration approach has been used to decrease the artefacts generated from the asymmetry in the substitution rates observed in the duplicates (Zhaxybayeval 2013). The inference was carried out using the Markov Chain Monte Carlo (MCMC) model with 10,000 samples in each 1,000 chains (Drummond et al. 2012). Subsequently, the first 1,000 samples were removed with the *burn-in* option in TreeAnnotator (Drummond et al. 2006), and then the estimated consensus tree was created with the best posterior probability (PP) for each node. The tree was visualized and customized with Figtree 1.4 (Rambaut 2009).

### Codon usage bias

Taking into account that codon usage bias may result in phylogenetic artefacts in gene trees (Inagaki et al. 2004, Inagaki and Roger 2006; Liu et al. 2014), due to differences in codon usage in the *saltans* and *willistoni* radiations in comparison to other *Drosophila* groups (Powell et al. 2003; Vicario et al. 2007) and because the *D. willistoni* phylogenetic position is commonly an artefact (Pélandakis and Solignac 1993; Tarrío et al. 2001; Gailey et al. 2000), the analyses were performed to estimate the *relative synonymous codon usage* (RSCU) by group and by gene. The RSCU was carried out with MEGA X (Kumar et al. 2018), along with CAIcal (Puigbò et al. 2008), to identify the number of effective codons (ENC) and the GC proportion at the third codon position (%GC3). We carried out a PCA to investigate the difference between the RSCU of Drosophilidae groups and a t test to verify the difference between the ENC and %GC3 between the clade *willistoni*-*saltans*-*Lordiphosa* and the rest of the Drosophilidae phylogeny. The statistical analyses were conducted in R v. 4.1.2 (R core team 2021).

### Relative rate of nucleotide substitution

To identify whether *unc-4* and its duplicates are evolving at different rates, the relative rate test was performed with PHYLTEST 2.0 (Kumar 1996). The external groups used were the *unc-4* sequences annotated from Steganinae species, applying Kimura 2-parameters (Kimura 1980) as the best substitution model.

### Estimates of selection pressure and investigation of signatures of positive selection

To characterize the selection acting on the *unc-4* and *OdsH* genes, codon-based likelihood methods were run using the CODEML package in PAML ver 4.9 (Yang 2007). Maximum likelihood estimates of the selection pressure were measured by the nucleotide substitution rate (ω = Ka/Ks) of nonsynonymous (Ka) to synonymous (Ks) substitutions. For these analyses, two trees in Newick format were used, one of which was *Tree 1*, described above, using the alignment of the sequences *unc-4* and *OdsH*. Since only the *OdsH* sequences presented nonsynonymous substitutions, selection tests were also performed considering only this gene, constructing a tree, hereafter referred to as *Tree 2*, also by Bayesian inference, with the same priors as for *Tree 1*, using only sequences (48) from groups of Drosophilinae represented by more than two species. For these analyses, the branch model test allows the ω ratio to vary among branches in the phylogeny and is useful for detecting positive selection acting on particular lineages (Yang 1998; Yang and Nielsen 1998).

Two types of branch model tests were used. Type 1 fits the so-called free-ratio model that assumes an independent ω ratio for each branch. Although this approach is very parameter-rich and its use is discouraged (Yang 2007), it was used to estimate ω in specific branches candidate for positive selection (ω >1) or relaxed negative selection (ω>0.3) using *Tree 2*. Type 2 allows different branch groups to have different ω, leading to so-called “two ratios”, by using branch labels in the tree file. This approach was applied to estimate the ω value in *Tree 1*, with labels in *unc-4* and *OdsH* ancestor and internal nodes, and in *Tree 2*, labelling each group of species and clades that presented signatures of relaxed negative selection in Model 1. All the hypotheses developed to identify the ω value were tested using the χ² test, with the comparison of the lnL values of each hypothesis.

### Transcription factor-binding sites at the *unc-4* and *OdsH* regulatory regions

To investigate the presence of different transcription factor-binding sites (TFBSs) located at the *unc-4* and *OdsH* regulatory regions, the sequences were extracted 500 bp upstream and downstream of the genes from all species in which expression had been analysed previously. In addition, *D. sechellia*, *D. simulans* and *D. mauritiana* were included because in these species, *OdsH* is associated with hybrid sterility. For this analysis, the *OdsH* regulatory sequences were subjected to enrichment analysis with CiiiDER (Gearing et al. 2019) to identify differentially enriched TFBSs between *unc-4* and *OdsH* by using the *unc-4* sequences as background. We used the JASPAR CORE (Castro-Mondagon et al. 2022) database of insect TFBSs for this analysis. The *deficit threshold* default (0.15) and the Fisher *p value* threshold 0.05 were applied. The transcription factors with differential enrichment of binding sites to the regulatory regions between *unc-4* and *OdsH* were used for Gene Ontology (GO) analysis (Ashburner et al. 2000; Mi et al. 2019) in the biological process category.

### Protein Functional Motifs

The homeodomains and the octapeptide were found in Unc-4 and OdsH proteins separately with MEME (Bailey and Elkan 1994) in the MEME Suite platform (Bailey et al. 2015). To observe the wide pattern of homeodomain diversity in both proteins from Drosophilinae, they were calculated with the translated sequences retrieved from the Drosophilinae alignment. The octapeptide was estimated from the alignment of the 11 C-terminal amino acids of the Unc-4 and OdsH proteins, as reported in NCBI (Supplementary Table S1, Supplementary Material online).

The binding stability of the tri-dimensional models for the Unc-4 and OdsH homeodomains associated with the DNA was assessed to infer whether their protein sequence divergence causes functional divergence in biophysical terms. The protein modelling of the Unc-4 and OdsH homeodomains was developed with SwissModel (Waterhouse et al. 2018) using the structure of PDB 3LNQ (Miyazono et al. 2010) as a template. The modelling was performed for *D. melanogaster* (NP_573242.2 and NP_523389.3) and for *T. dalmani* Unc-4 (XP_037943702.1) as an outgroup to the duplication event. Afterwards, the complexes derived from the structural model Unc-4 from *T. dalmani* and the DNA structure were minimized from molecular dynamic simulations using GROMACS (Abraham et al. 2015), applying the AMBER14-OL15 package with ff14sb protein (Maier et al. 2015) and ff99bsc0OL15 DNA (Zgarbová et al. 2015) force fields, as well as the TIP3P1 water model (Jorgensen et al. 1983).

The molecular system was inserted into a solvated cubical box by a 100 mM NaCl solution in water. Energy minimization was performed with the *steepest descent integrator* and the conjugated gradient algorithm, with 500 kJ·mol^−1^·nm^−1^ as the maximum force threshold. The calculation of the perturbation values of the variation in the free energy of ligation (ΔΔGb) was assessed with the observed OdsH substitutions in *Drosophila*, which interferes with the stability of the homeodomain/DNA complex, by using the mCSM server (Pires et al. 2014), in comparison to the Unc-4/DNA homeodomain complex structure.

### *OdsH* and *unc-4* Expression

The expression profiles of *unc-4* and *OdsH* were manually inspected with the *Tracks* tool from the *Gene* platform available at NCBI (www.ncbi.nlm.nih.gov/gene) using public databases (*D. pseudoobscura*: PRJNA291085; *D. grimshawi*: PRJNA317989: *T. dalmani*: PRJNA240197; other species: PRJNA388952). All Drosophilinae species with available transcriptome expression data from either reproductive or nonreproductive tissues were analysed separately by sex. The same approach was used to identify the expression of the single-copy *unc-4* gene in the *Teleopsis dalmani* genome as an outgroup to the duplication event. For each species and tissue, the genes were characterized as expressed when they had > 10 *counts* identified at the expression histogram from the *Tracks* tool.

Experimental analysis of *OdsH* expression was conducted in *D. mojavensis baja* and *D. arizonae* and their hybrids. For this, intra- and interspecific crosses were performed in both directions between *D. arizonae* from Metztitlan, Hidalgo, Mexico (Stock Center n.: 15081-1271.**17**), and *D. m. baja* from the Cape Region, Santiago, Baja California Sur, Mexico (Stock Center n.: 15081-1352.**20**). These species were chosen as representatives of the *Drosophila* subgenus, allowing the observation of *OdsH* functions outside the *Sophophora* subgenus previously reported. In addition, they show recent divergence and incomplete reproductive isolation. Their reciprocal interspecific crosses are asymmetrical, with the male offspring being fertile when descended from male *D. arizonae* (H♀moj^baja^♂ari) and sterile when descended from male *D. m. baja* (H♀ari♂moj^baja^) and the female offspring being fertile in both directions (Banho et al. 2021). Since deregulation in hybrids might result from fast male evolution, the comparison between fertile and sterile hybrids can help to determine specific deregulation related to sterility (Gomes and Civetta 2014, 2015). For the experimental crosses, virgin males and females were collected until 48 h after emergence and isolated in tubes containing *Opuntia* sp.-based media for 3 days. For this, each cross was performed in 35 replicates, each containing 10 couples, for 12 days. The testes of descendants (10-12 days) were dissected in 1X PBS.

Dissected testes in 1X PBS from both hybrids and parental species were subjected to smRNA FISH to determine if *OdsH* had atypical expression in sterile hybrids, considering the spermatogenesis phases. The testes were then fixed in fixing buffer 4% formaldehyde, 0.3% Trirton X-100, 1x PBS) for 20 min at room temperature, rinsed three times in 0.3% Triton X-10, one in PBS, and permeabilized in 70% ethanol at 4°C overnight. Permeabilized testes were rehydrated in smRNA FISH wash buffer (10% formamide in 2x SSC) for 10 min.

Testes were resuspended in 50μL hybridization buffer (10% dextran sulfate, 10% formamide in 2x SSC, supplemented with 1μL of smRNA FISH probes) designed with Stellaris® Probe Designer version 4.2 (https://www.biosearchtech.com/stellaris-designer) (supplementary table S8, Supplementary Material online), synthesized and labeled with ATTO 550. Hybridization was performed with rotation at 37°C overnight. Testes were then washed twice with smRNA FISH wash buffer at 37°C for 30 min and twice with 2x SSC solution. Then, DNA was stained with DAPI (Thermo Fisher Scientific) (1/500 dilution in 2x SSC) at room temperature for 20 min. Images were captured using an upright Zeiss LSM780-NLO confocal microscope.

For quantitative analysis, the RNA was extracted from the testes of seven biological replicates each using 25 individuals using the RNeasy kit (Qiagen) and was treated with DNase (DNA-free kit; Ambion). For each replicate, 1000 ng of RNA was converted to cDNA using a *High Capacity cDNA Reverse Transcription kit* (Thermo Fisher). The relative level of mRNA was quantified using specific oligonucleotides and probes (TaqMan, Thermo Fisher Scientific) for *OdsH* (forwards primer: AGCCGCAGAGCTGCA; reverse primer: GCTCGATCGCCTTGGCTAT; probe: CTGCAGGAGCTGCGAGCCA). qPCR was then conducted using three technical replicates, each containing 100 ng of cDNA in a *LightCycler* 480 (Roche Diagnostics). The expression level was measured by the RQ ratio in relation to the *endogenous ribosomal gene 49* (*rp49*), also known as nrpL (forwards primer: CCCAACATTGGTTACGGTTCCA; reverse primer: GCACATTGTGTACGAGGAATTTCTT; probe: CACCCGCCACATGCT). Then, the relative quantity of the transcripts was normalized by the expression (RQ = E^Ct^ ^rp49^/E ^Ct^ ^OdsH^; E = reaction efficiency). The normalized values were subjected to Shapiro‒Wilk and Bartlett tests for each tissue. Since they did not present a normal distribution and variance homogeneity, their variances were calculated through the Kruskal‒Wallis test.

## Supplementary Material

Supplementary data are available in supplementary information files, Supplementary Material Online.

## Data Availability

All data generated in this study are included in supplementary information files, Supplementary Material Online.

## Acknowledgments

We acknowledge funding from The São Paulo Research Foundation (FAPESP)/ Université de Lyon 1 (UdL – IDEXLyon) Joint Call to C.M.A.C [2020/06238-2] and to CV, from the National Council for Scientific and Technological Development (CNPq) to C.M.A.C [308020/2021-9] and from the International fellowship from UdL – IDEXLyon – FAPESP to WVBN. This study was financed in part by the Coordenação de Aperfeiçoamento de Pessoal de Nível Superior – Brasil (CAPES) – Finance Code 001 to WVBN.

## Author Contributions

W.V.B.N. and C.M.A.C. designed the study. W.V.B.N., D.S.O., G.R.D., A.B.C., I.C.P., J.M.B., N.G., A.A., N.B., C.V. and C.M.A.C. collected and/or analysed data. W.V.B.N., C.V. and C.M.A.C. wrote the manuscript. W.V.B.N., D.S.O., G.R.D., A.B.C., I.C.P., J.M.B., N.G., A.A., N.B., C.V. and C.M.A.C. reviewed and edited the manuscript. All authors contributed to the article and approved the submitted version.

## Conflict of Interest

We declare there are no conflicts of interest.

## Notes

### Competing Interest Statement

The authors have declared no competing interest.

