## Supplementary figures for "A comprehensive evolutionary scenario for the origin and neofunctionalization of the *Drosophila* speciation gene *Odysseus* (*OdsH*)"

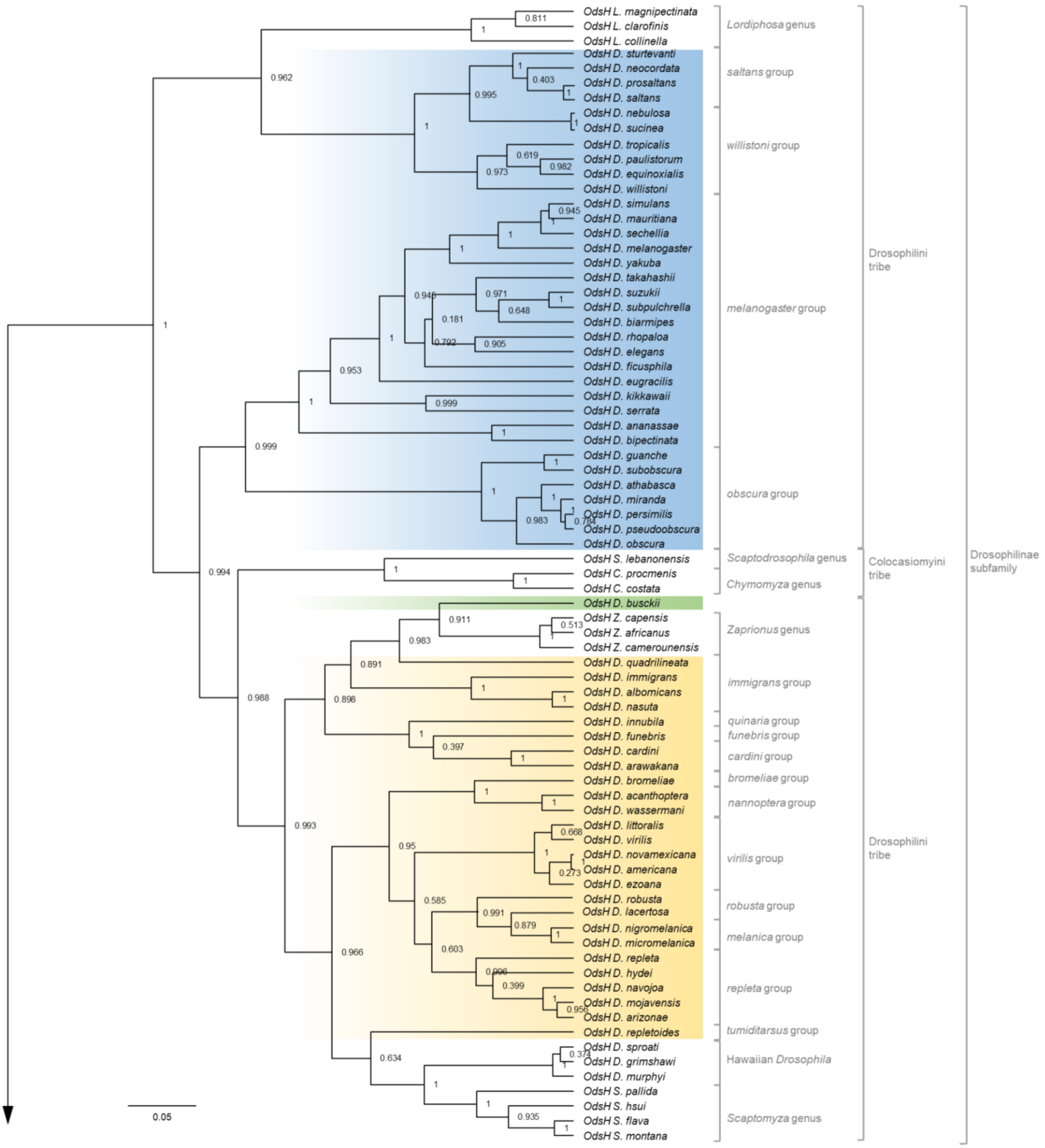

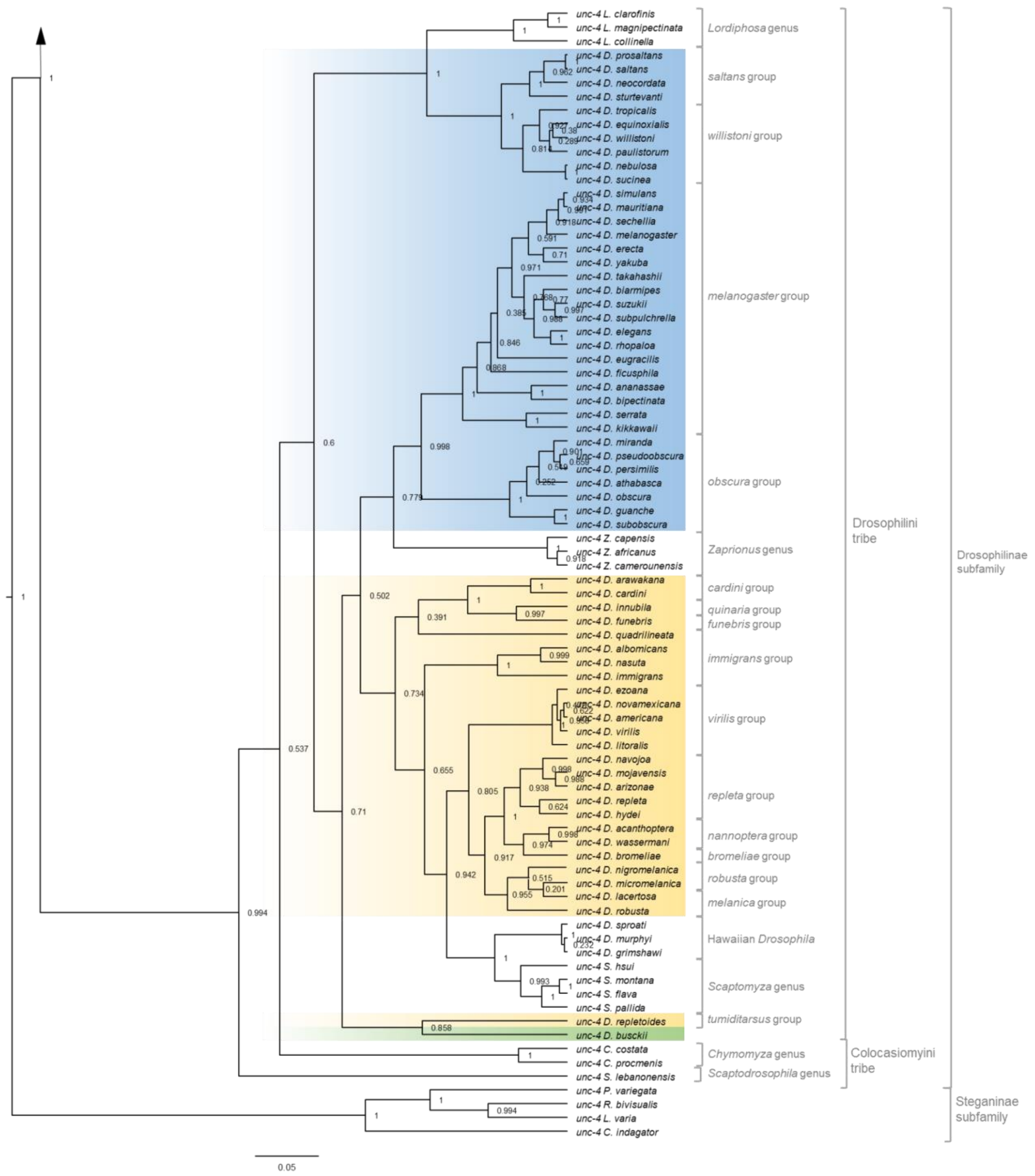


**Supplementary Fig. S1**. Bayesian inference of the phylogenetic relationships between the sequences of the paralogue genes *unc-4* and *OdsH*, using the GTR+G+I model of nucleotide substitution. The analysis was performed with 405 sites from 153 nucleotide sequences. All positions containing gaps and ambiguous positions were removed from the pairwise sequence analysis. At the root of each clade, the posterior probabilities are indicated. The analysis was conducted in BEAST v16.1. The *unc-4* clade, subdivided into more basal single copy *Steganinae* (outgroup) and Drosophilinae, is presented at the base of the phylogeny followed by the *OdsH* clade in the upper part. Arrows indicate the junction between the two parts of the phylogeny. Subgenera are highlighted in blue (*Sophophora*), yellow (*Drosophila*) and green (*Dorsilopha*). The brackets indicate *Drosophila* species groups and genera, tribes and subfamilies of Drosophilidae.

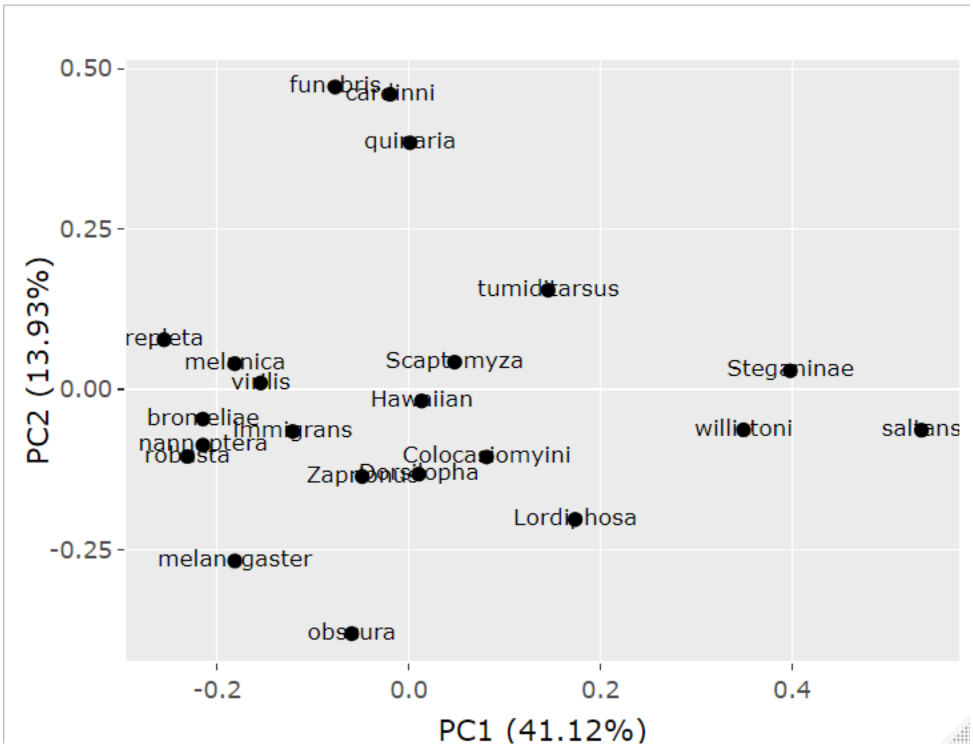


**Supplementary Fig. S2**. PCA of the codon usage estimated by the RSCU calculation of *unc-’* sequences of Drosophilinae.

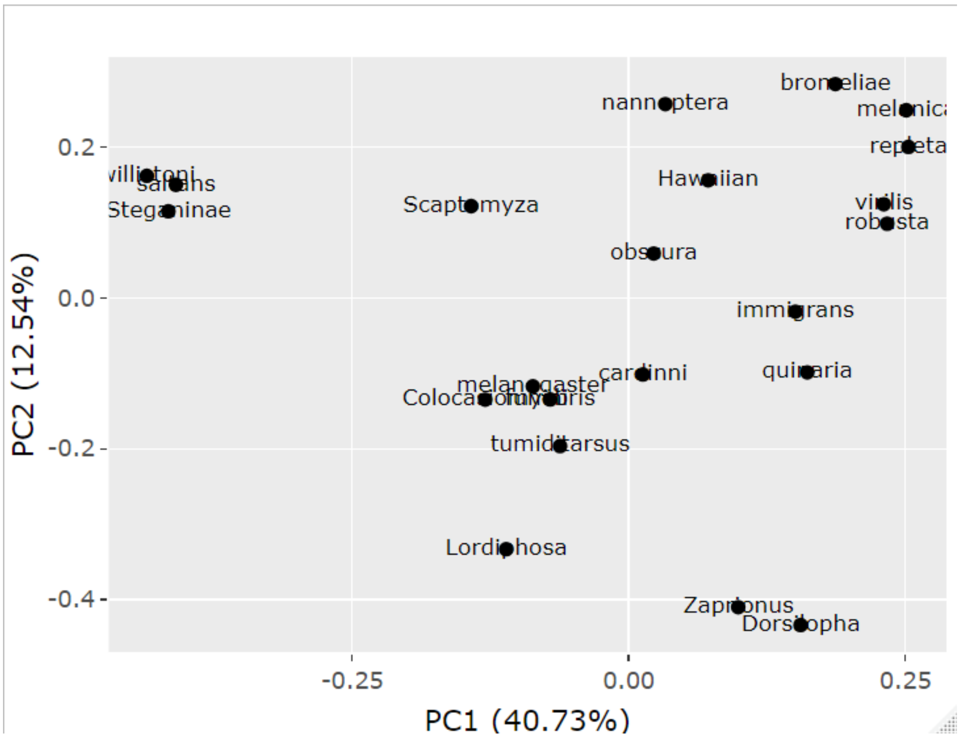


**Supplementary Fig. S3**. PCA of the codon usage estimated by the RSCU calculation of *OdsH* sequences of Drosophilinae and the orthologue single copy *unc-4* in Steganinae.

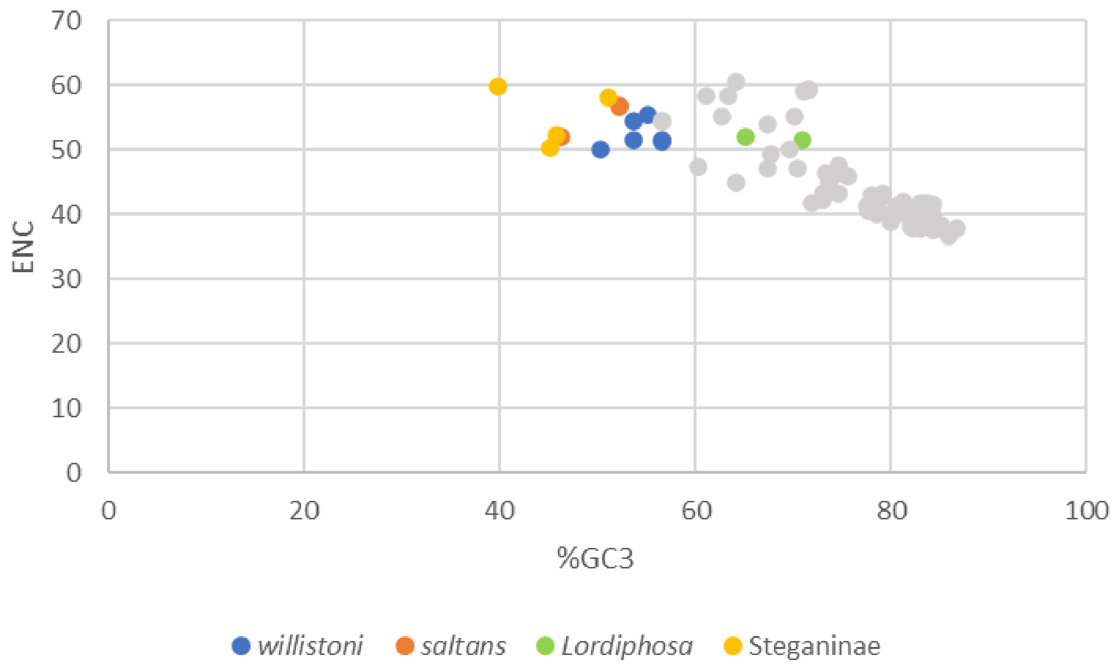


**Supplementary Fig. S4**. % of GC3 and ENC in Drosophilidae *unc-4* sequences.

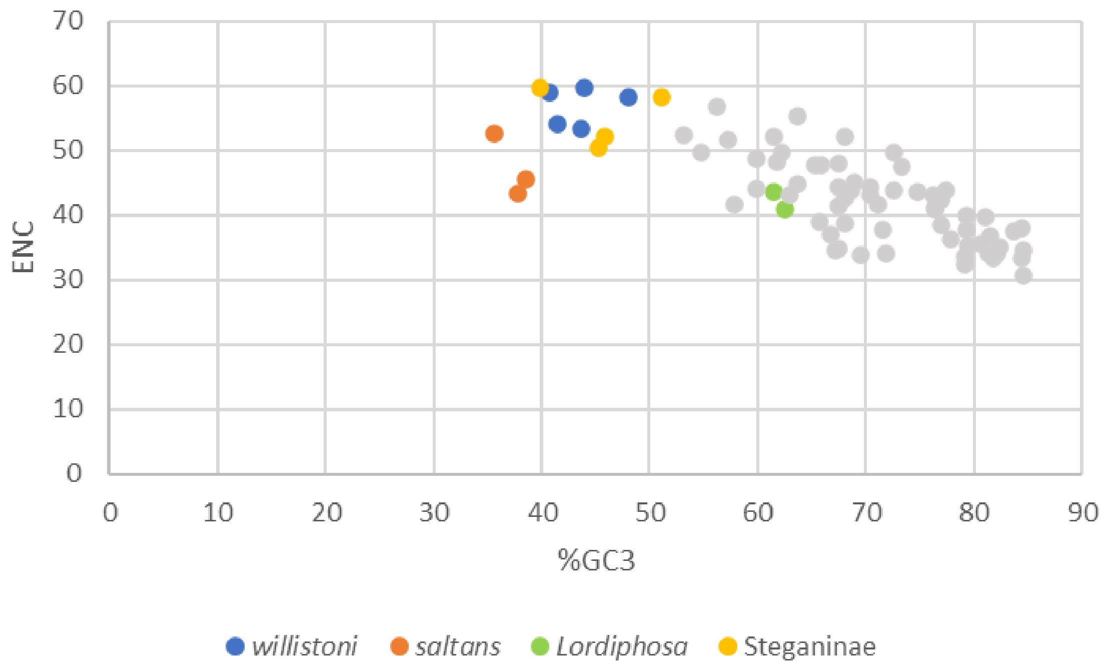


**Supplementary Fig. S5**. % of GC3 and ENC in *OdsH* sequences of Drosophilinae and the orthologue single copy *unc-4* in Steganinae.

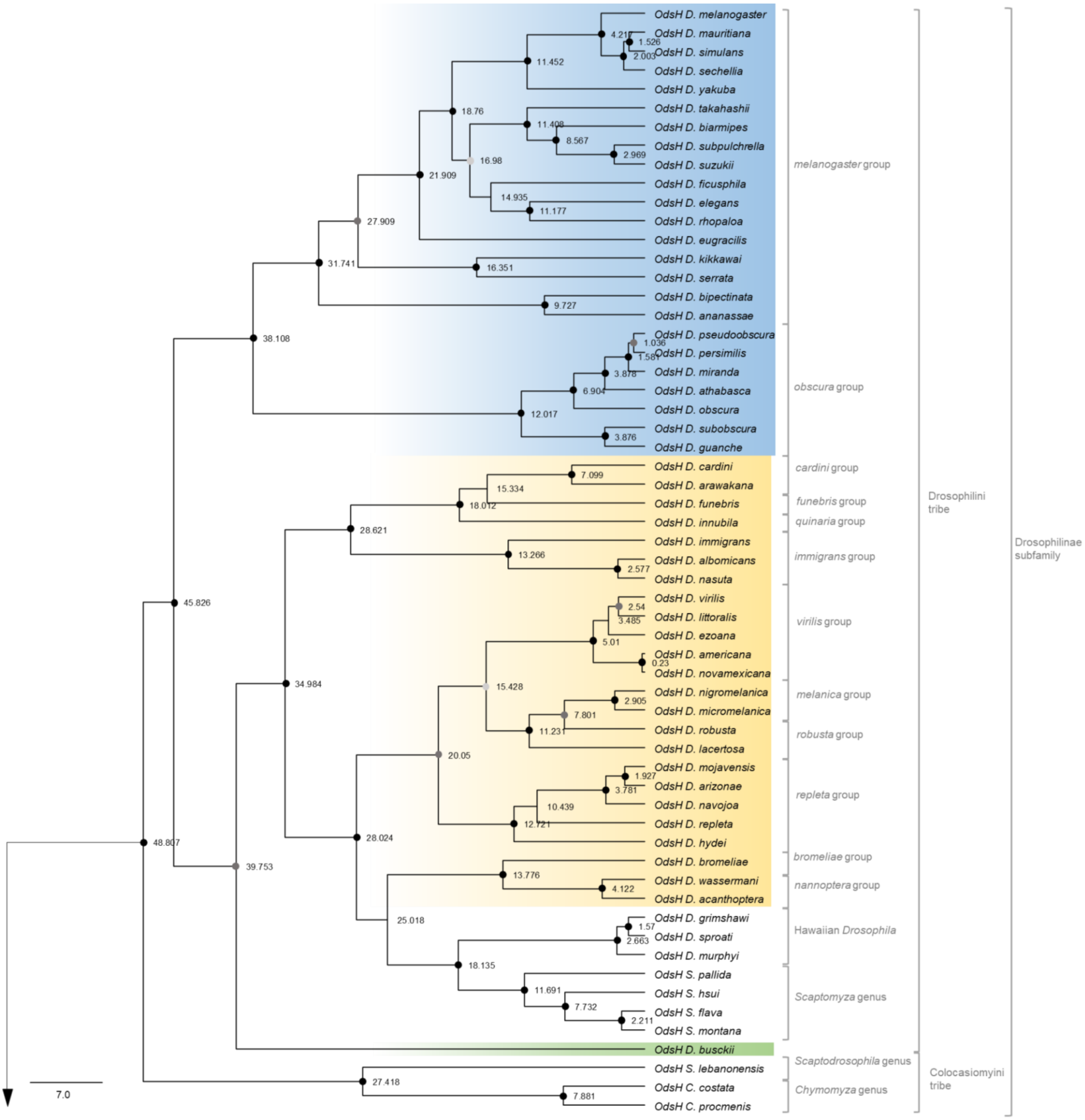
OdsH D. melanogaster
OdsH D. mauritiana
OdsH D. simulans
OdsH D. sechellia
OdsH D. yakuba
OdsH D. takahashii
OdsH D. biarmipes
OdsH D. subpulchrella
OdsH D. suzukii
OdsH D. ficusphila
OdsH D. elegans
OdsH D. rhopaloa
OdsH D. eugracilis
OdsH D. kikkawai
OdsH D. serrata
OdsH D. bipectinata
OdsH D. ananassae
OdsH D. pseudoobscura
OdsH D. persimilis
OdsH D. miranda
OdsH D. athabasca
OdsH D. obscura
OdsH D. subobscura
OdsH D. guanche
OdsH D. cardini
OdsH D. arawakana
OdsH D. funebris
OdsH D. innubila
OdsH D. immigrans
OdsH D. albomicans
OdsH D. nasuta
OdsH D. virilis
OdsH D. littoralis
OdsH D. ezoana
OdsH D. americana
OdsH D. novamexicana
OdsH D. nigromelanica
OdsH D. micromelanica
OdsH D. robusta
OdsH D. lacertosa
OdsH D. mojavensis
OdsH D. arizonae
OdsH D. navojoa
OdsH D. repleta
OdsH D. hydei
OdsH D. bromeliae
OdsH D. wassermani
OdsH D. acanthoptera
OdsH D. grimshawi
OdsH D. sproati
OdsH D. murphyi
OdsH S. pallida
OdsH S. hsui
OdsH S. flava
OdsH S. montana
OdsH D. busckii
OdsH S. lebanonensis
OdsH C. costata
OdsH C. procmenis
melanogaster group
obscura group
cardini group
funebris group
quinaria group
immigrans group
virilis group
melanica group
robusta group
repleta group
bromeliae group
nannoptera group
Hawaiian Drosophila
Scaptomyza genus
Scaptodrosophila genus
Chymomyza genus
Drosophilini tribe
Colocasiomyini tribe
Drosophilinae subfamily
7.0

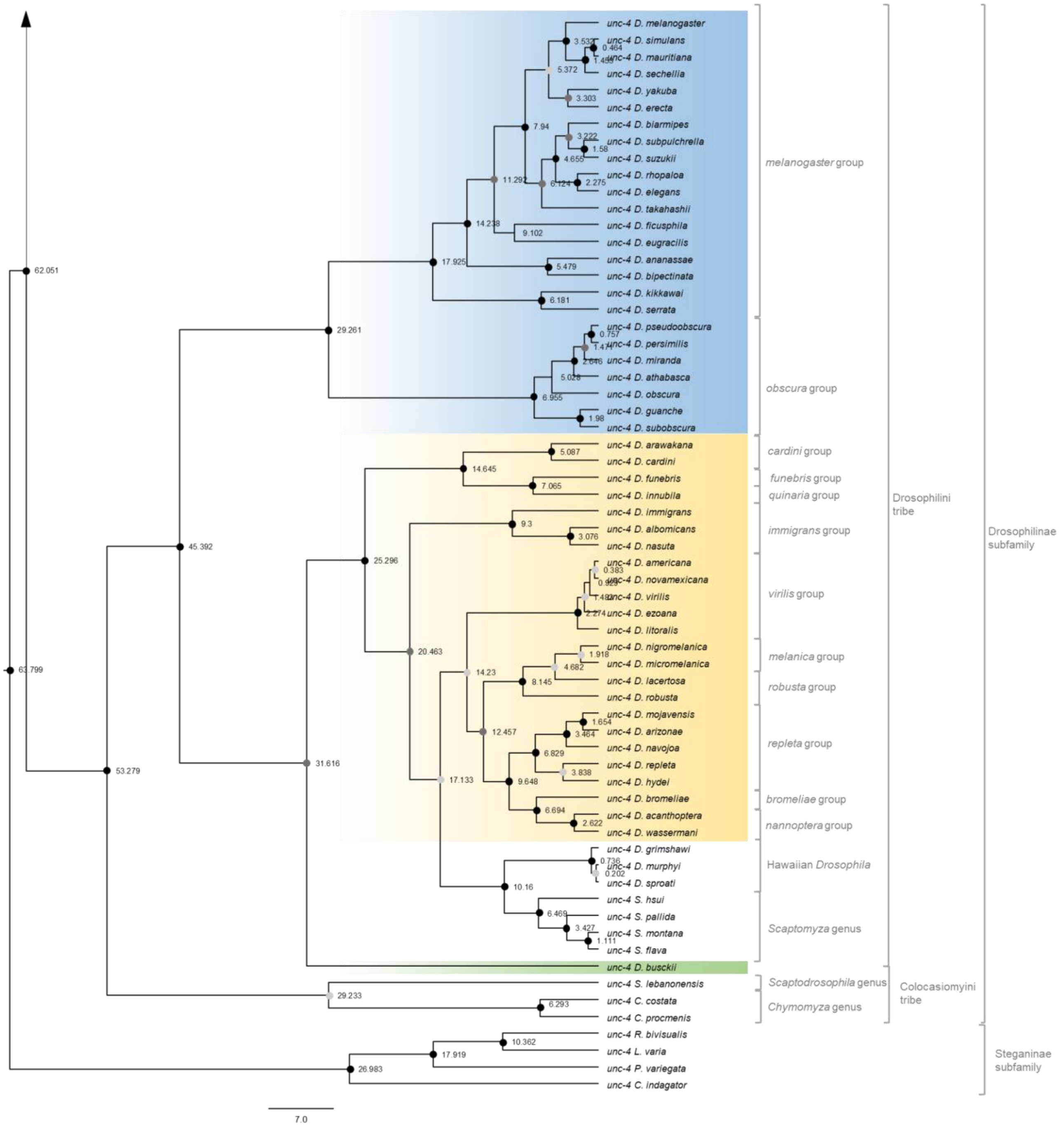


**Supplementary Fig. S6.** Calibrated Bayesian phylogenetic inference of the sequences of the paralogue genes *unc-4* and *OdsH*, using the GTR+G+I model of nucleotide substitution. The analysis was performed with 405 nucleotide sites from 131 nucleotide sequences. All positions containing gaps and ambiguous bases were removed from the pairwise sequence analysis. The branches referring to the *Drosophila* taxonomic groups were compressed. At the root of each clade, the posterior probability is presented by black (>0.9) and grey (>0.7) circles and the estimated time of divergence are indicated. The analysis was conducted in BEAST v16.1. The *unc-4* clade, subdivided into more basal single copy *Steganinae* (outgroup) and Drosophilinae, is presented at the base of the phylogeny followed by the *OdsH* clade in the upper part. Arrows indicate the junction between the two parts of the phylogeny. Subgenera are highlighted in blue (*Sophophora*), yellow (*Drosophila*) and green (*Dorsilopha*). The brackets indicate *Drosophila* species groups and genera, tribes and subfamilies of Drosophilidae. The clade *willistoni-saltans-Lordiphosa* was not included in this analysis.

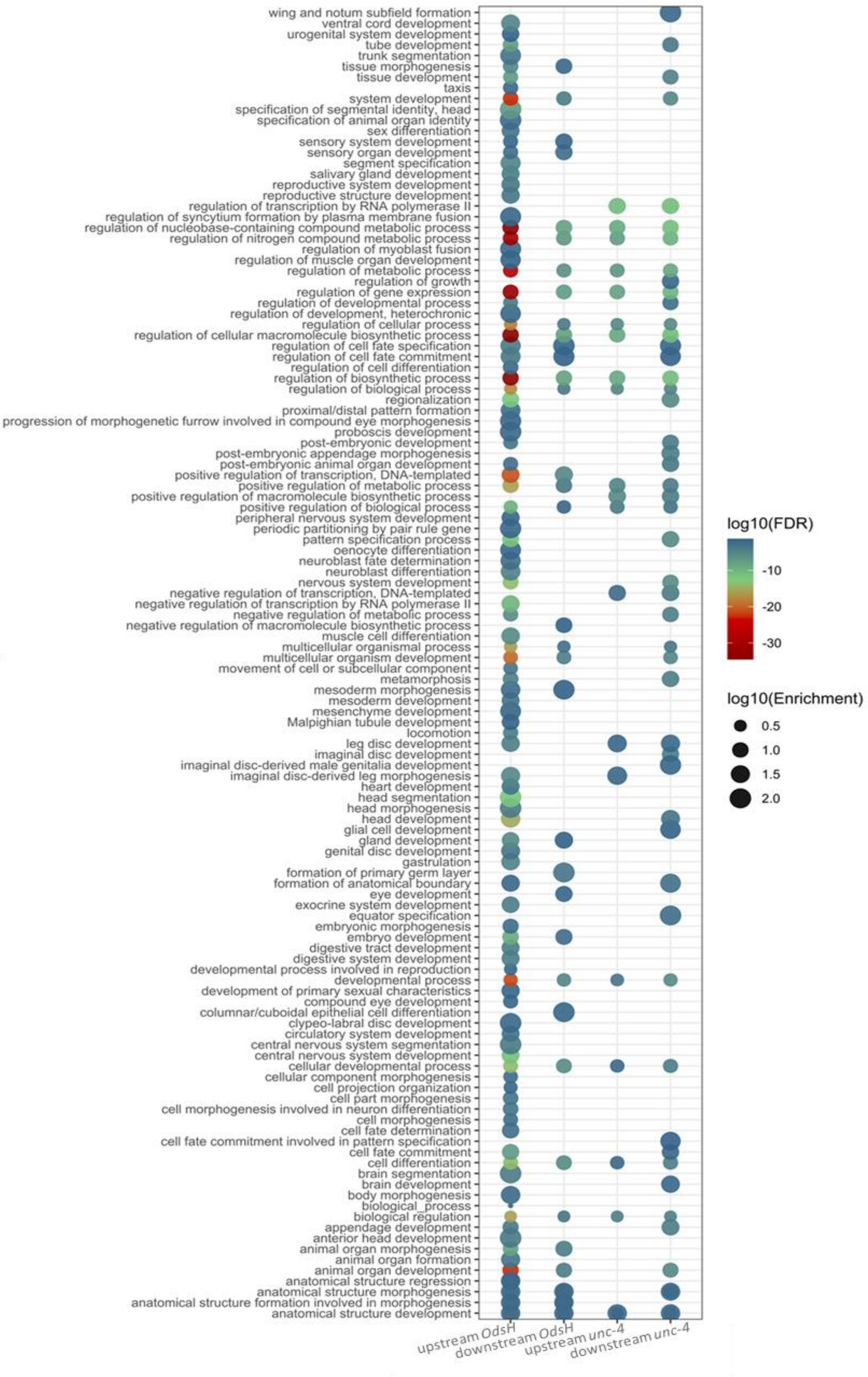


**Supplementary Fig. S7.** Enriched GO terms for biological process category of TFBS in the regulatory region of *OdsH* and *unc-4* of *Drosophila.*

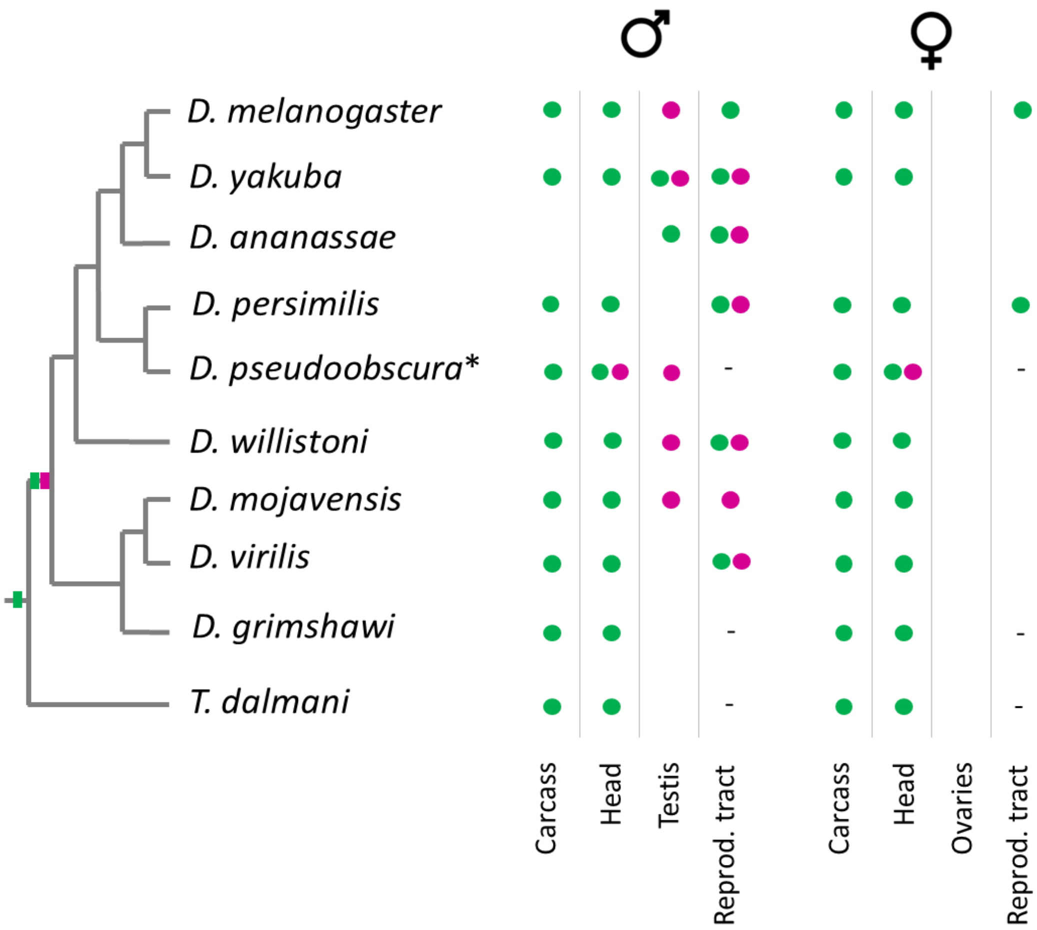


**Supplementary Fig. S8.** Expression of *OdsH* (pink) and *unc-4* (green) in each tissue of *Drosophila.* The representation of the phylogenetic relationships of the species was based on Suvorov et al. (2022). The squares on phylogeny represent the presence of the genes *OdsH* (pink) and *unc-4* (green). *Teleopsis dalmani* is the outgroup that presents *unc-4* single copy.

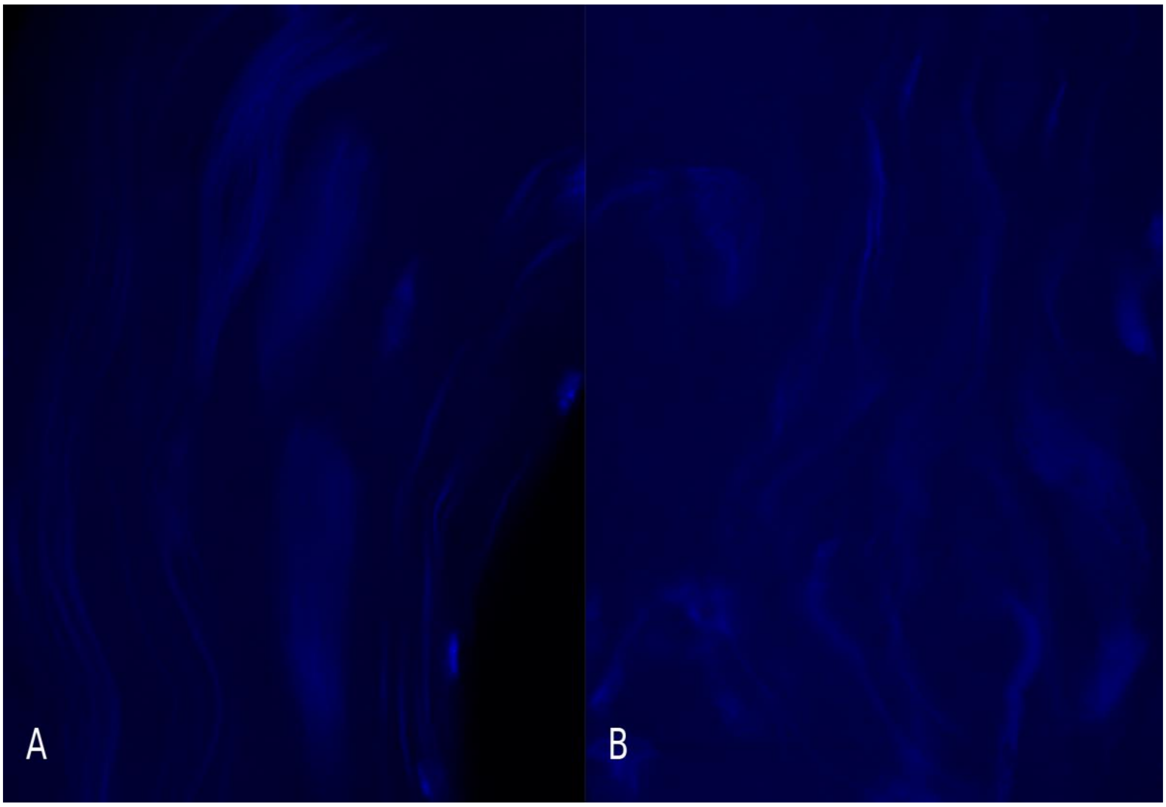


**Supplementary Fig. S9.** Sperm bundles in hybrids from *D. m. baja* and *D. arizonae*. A. H♀moj$^{baja}$♂ari (fertile). B. H♀ari♂moj$^{baja}$ (sterile). NOTE: DAPI (blue) was used to label DNA.

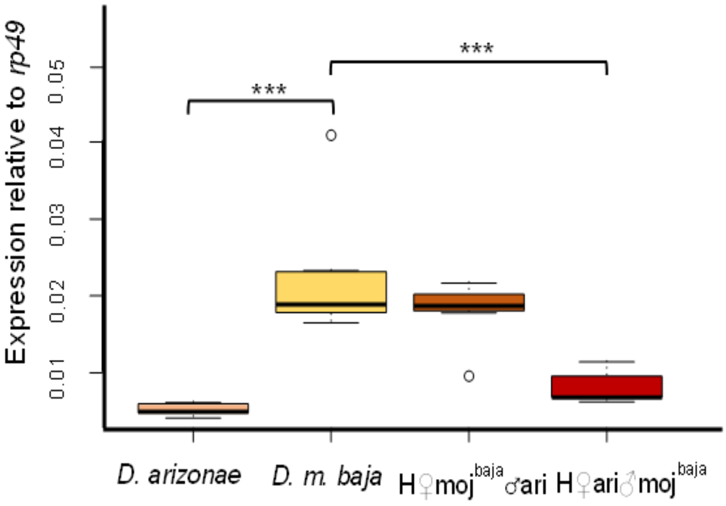


**Supplementary Fig. S10.** Relative expression of *OdsH*, in relation to *rp49*, in tissues *D. arizonae* and *D. m. baja* and their reciprocal hybrids. NOTE: *** $p<0.01$
